# Age-dependent miR156-targeted *SPLs* are required for extrafloral nectary development in *Passiflora* spp

**DOI:** 10.1101/2024.02.20.581215

**Authors:** Jéssica Ribeiro Soares, Kerly Jessenia Moncaleano Robledo, Vinicius Carius de Souza, Lana Laene Lima Dias, Lázara Aline Simões Silva, Emerson Campos da Silveira, Claudinei da Silva Souza, Elisandra Silva Sousa, Pedro Alexandre Sodrzeieski, Yoan Camilo Guzman Sarmiento, Elyabe Monteiro de Matos, Thais Castilho de Arruda Falcão, Lilian da Silva Fialho, Valéria Monteze Guimarães, Lyderson Facio Viccini, Flaviani Gabriela Pierdona, Elisson Romanel, Jim Fouracre, Wagner Campos Otoni, Fabio Tebaldi Silveira Nogueira

## Abstract

- Passion flower extrafloral nectaries (EFNs) protrude from adult leaves and facilitate mutualistic interactions with insects, but how age cues control EFN establishment remains poorly understood.
- Here, we combined genetic and molecular studies to investigate how leaf development and EFN patterning are regulated through the age-dependent miR156-*SQUAMOSA PROMOTER BINDING PROTEIN LIKE* (*SPL*) module in two EFN-containing *Passiflora* species with distinct leaf shapes.
- Low levels of miR156 correlate with leaf maturation and EFN formation in *Passiflora edulis and P. cincinnata*. Consistently, overexpression of miR156 (miR156-OE), which leads to low levels of *SPLs*, affected leaf ontogeny and EFN development in both species. Laminar EFNs were underdeveloped and less abundant in both *P. edulis and P. cincinnata* miR156-OE leaves. Importantly, the ecological relationships established by EFNs and their sugar profiles were negatively regulated by high levels of miR156. Moreover, transcriptome analysis of young leaf primordia revealed that miR156-targeted *SPLs* may be required for proper expression of leaf and EFN development-associated genes in *P. edulis and P. cincinnata*.
- Our work provides the first evidence that the highly conserved miR156/*SPL* module regulates EFN development in an age-dependent manner and that the program responsible for EFN development is closely associated with the heteroblastic developmental program of the EFN-bearing leaves.

## Introduction

In response to endogenous and environmental cues, plants undergo a series of developmental transitions during their life cycles, including the switch from the juvenile to the adult phase of vegetative growth (known as vegetative phase change). Depending on the species, vegetative phase change can be accompanied by shifts in a wide variety of traits, including the formation of secretory structures (Leichty & Poethig, 2019). In many plant species, the development or increase in the abundance of extrafloral nectaries (EFNs) occurs as the plant ages (Villamil et al., 2013; Moraes et al., 2022; Kwok & Laird, 2012; Leichty & Poethig, 2019). For example, the abundance and complexity of EFNs increase in *Turnera velutina* as the plant progresses into the reproductive phase (Villamil et al., 2013). In *Passiflora organensis,* EFNs are absent in the juvenile vegetative stage, with the adult stage characterized by EFN formation on the abaxial side of the leaf lamina (Moraes et al., 2022).

As secretory structures, nectaries secrete a chemically complex compound called nectar, which is rich in carbohydrates but can also contain amino acids, proteins, alkaloids and phenolics (Heil, 2011; Marazzi et al., 2013; Cardoso-Gustavson et al., 2013). Floral nectaries (FNs) are found in the floral whorls and, together with floral morphology, support a co-evolutionary relationship with pollinators (Ulmer & MacDougal, 2004; Pérez & d’Eeckenbrugge, 2017). On the other hand, EFNs are located on leaves, leaf petioles, stipules or floral bracts and play a role in the indirect defense of plants by attracting insects, mostly ants, which protect the plant from herbivore attacks in exchange for nectar, establishing a mutualistic interaction (Elias, 1983; Apple & Feener, 2001; Heil, 2015). EFNs have a plethora of effects reflected in the increased longevity and efficiency of entomophagous insects that consume nectar as an energy source, and increased crop productivity by contributing to reduced herbivory (Heil, 2015).

Recent investigations into the genetic basis of nectary development have provided evidence that FNs and EFNs share developmental genetic networks (Lee et al., 2005; Pei et al., 2021). However, it is important to consider that EFN development may recruit different regulatory pathways than those required for FNs (Marazzi et al., 2013). Nectary development is thought to be regulated by the same genetic program that is present and active in the organ that exhibits this secretory structure (Lee et al., 2005; Marazzi et al., 2013; Leichty & Poethig, 2019). For example, changes in the morphology of EFNs present in the main vein of leaves of *Gossypium arboreum* have been shown to accompany leaf ontogeny (Hu et al., 2020). Although the YABBY family gene *CRABS CLAW* (*CRC*) has been recognized as an important regulator of FN development in distinct species such as *G. hirsutum* and *Petunia hybrida* (Lee et al., 2005), the underlying mechanisms regulating the development and growth of EFNs are poorly understood, as most genetic and molecular studies have focused on FNs.

The genus *Passiflora* is represented by more than 500 species distributed in four subgenera: *Passiflora*, *Decaloba*, *Astrophaea*, and *Deidamioides*, with the subgenus *Passiflora* being the most numerous of the genus (Feuillet & MacDougal, 2007; Pérez & d’Eeckenbrugge, 2017). Floral and extrafloral nectaries are common in many *Passiflora* species, making the genus relevant for studies on the evolution, development and ecology of nectaries and nectar metabolites (Apple & Feener, 2001; Cardoso-Gustavson et al., 2013; Rocha et al., 2009; Pérez & d’Eeckenbrugge, 2017; Moraes et al., 2022). Comparative analyses of EFN patterning suggest a shared developmental program with leaf development to create EFN diversity in *Passiflora* species. Importantly, the final location and morphology of EFNs depends on the developmental program active in the organ that bears them (Krosnick et al., 2011).

*Passiflora edulis* and *P. cincinnata* are perennial climbing vines, in which EFNs are located adaxially in the petiole (petiolar EFN) and abaxially on the leaf margin (laminar EFN). Juvenile *P. edulis* plants produce monolobed leaves whereas leaves produced during the adult vegetative phase are trilobed. Petiolar EFNs in *P. edulis* are present on both juvenile and adult leaves, whilst laminar EFNs are observed only on adult leaves (Silva et al., 2019). In contrast, *P. cincinnata* petiolar and laminar EFNs are observed in trilobed juvenile leaves, and exhibit a similar distribution in the adult phase, when the leaves become five-lobed. These gradual changes in leaf patterning during the transition from juvenile to adult phase are known as leaf heteroblasty and have been documented for many species in the genus *Passiflora* (Chitwood & Otoni, 2017 a, b; Silva et al., 2019).

The microRNA156 (miR156) is the master regulator of vegetative phase change. The expression levels of miR156 decrease during juvenile development, leading to an increase in the activity of its targets in the *SQUAMOSA PROMOTER-BINDING PROTEIN-LIKE* (*SPL*) family of transcription factors. miR156-targeted *SPL* genes promote adult vegetative traits and therefore the timing of vegetative phase change is determined by the balance of miR156 and *SPL* transcript levels (Wu et al., 2009; Xu et al., 2016; He et al., 2018; Silva et al., 2019; Poethig & Fouracre, 2024). The miR156-*SPL* module is highly conserved among angiosperms, and has been found to regulate multiple plant developmental processes, including leaf morphology and maturation, trichome development, sugar metabolism, axillary bud initiation, shoot architecture, fruit development, stress responses, plant defense and bixin biosynthesis (Wu et al., 2009; Yu et al., 2010; Poethig, 2013; Silva et al., 2014; Feng et al., 2016; Arshad et al., 2017; Guo et al., 2017; Silva et al., 2019; Lawrence et al., 2021; Machado et al., 2023; Barrera-Rojas et al., 2023; Ferigolo et al., 2023). Heteroblastic leaf development in *P. edulis* is correlated with the dynamic expression patterns of miR156 and *PeSPL9* (Silva et al., 2019), suggesting that the miR156-*SPL* module regulates shoot maturation in *Passiflora*. In addition, the age-dependent development of EFNs in *Vachellia* (Fabaceae) species is correlated with a decrease in miR156 transcripts and increased expression of miR156-targeted *SPLs* (Leichty & Poethig, 2019), suggesting that the miR156-*SPL* module may have been widely coopted into regulating the timing of EFN emergence. However, a functional interaction between miR156 activity and EFN production has yet to be demonstrated.

Here, we investigated whether leaf ontogeny modulates EFN development through the miR156-*SPL* module in two *Passiflora* species (*P. edulis* and *P. cincinnata*), which exhibit distinct leaf morphology (Chitwood & Otoni, 2017a). Low levels of miR156-targeted *SPLs* in miR156-overexpressing *P. edulis* and *P. cincinnata* plants prolonged the juvenile phase and affected heteroblastic leaf patterning. Most importantly, anatomical, transcriptome, and sugar content analyses revealed that miR156-targeted *SPLs* are required for proper *Passiflora* laminar EFN ontogeny and mutualistic interaction with ants. Our results indicate that the development of EFNs is modulated by a preexisting genetic program in the organ bearing them, and that the establlishment and abundance of laminar EFNs in *Passiflora* species is correlated with the formation of lobes during the age-dependent heteroblastic leaf development.

## Materials and Methods

### Generation of microRNA156 overexpression lines in *Passiflora cincinnata* Mast. and *Passiflora edulis* Sims.’

Seeds of *Passiflora cincinnata* Mast. cv. BRS ‘Sertão Forte’ and *Passiflora edulis* Sims. ‘FB-300’ were kindly donated by Dr. Fábio Gelape Faleiro (EMBRAPA Cerrados, Brazil) and by Viveiros Flora Brazil Ltda. (Araguari, MG, Brazil), respectively. Subsequently, seeds were mechanically decoated and were surface-sterilized in 70% ethanol, for 60 s, under a laminar flow hood. Following, immersed in a commercial sodium hypochlorite solution – 2.0-2.5% active chlorine plus two drops of Tween 80 (for each 100 mL of bleaching solution) for 15 min. The material was rinsed thrice (30 s duration each) with sterile water. Approximately ten decoated seeds were inoculated in an MS medium containing MS basal salts and vitamins (Murashige and Skoog, 1962) (PhytoTech^®^ Labs, Kansas), 0.01% (w/v) myo-inositol (Sigma Chem. Co., St. Louis), 3.0% (w/v) sucrose (PhytoTech^®^ Labs), and 0.6% (w/v) agar (Plant TC Micropropagation Grade; (PhytoTech^®^ Labs). The pH of the medium was adjusted to 5.8 ± 0.05, prior to the addition of the gelling agent. Seeds were kept for 15 days in the dark, at 25 ± 1 °C, followed by 15 days under a 16-h photoperiod and irradiance of 50 ± 3 μmol m^-2^ s^-1^ .

Genomic fragment harboring the *AtMIR156a* precursor was amplified from *Arabidopsis thaliana,* as described Feng et al. (2016)*. Agrobacterium tumefaciens* strain AGL-1 with a miR156 overexpression construct (p35S: HygR pUbi10-MIR156a-AtuOCS pMAS-CRT1) was cultivated in liquid LB medium containing 50 mg L^-1^ rifampicin (PhytoTech^®^ Labs) and 100 mg L^-1^ kanamycin (PhytoTech^®^ Labs), and carbenicillin (Gibco, Grand Island, NY) for 48 h at 28°C and 120 rpm.

*Passiflora* transformation protocol was done according to Manders et al. (1994) and Otoni et al. (2007), with few modifications. Thirty-d-old seedlings (15 days dark/15 days light) were used as a source of explants. Hypocotyls were sectioned transversally in segments of 0.8 – 1.0 cm in the suspension of *A. tumefaciens*, and after 15-20 min, the excess suspension was discarded. The explants were transferred to the regeneration medium composed of MS salts and vitamins supplemented with 3.0% (w/v) sucrose, 100 mg L^-1^ myo-inositol, 1 mg L^-1^ 6-benzylaminopurine (BA; PhytoTech Labs^®^) and 100 μM acetosyringone, starting the co-culture stage under dark conditions at 25°C, for two days. After, the explants were triple rinsed with liquid MS medium containing 300 mg L^-1^ of Timentin (PhytoTech^®^ Labs), and 250 mg L^-1^ Cefotaxime (PhytoTech Labs^®^). Subsequently, it was transferred to the semi-solid MS-based selective medium, with BA at 1 mg L^-1^, hygromycin (PhytoTech Labs^®^) at 6 mg L^-1^, Norflurazon (Sigma Chem. Co.) at 3.8 mg L^-1^, and Timentin at 300 mg L ^-1^. Explants were kept at 25 ± 1 °C under a photoperiod of 16:8 h and irradiance of 50 ± 3 μmol m^-2^ s^-1^ for six weeks, and every 15 days recultured to a freshly prepared selective culture. In the elongation phase, well-developed shoots (above 0.5 mm) were transferred to flasks containing hormone-free MS medium supplemented with hygromycin and Norflurazon.

### Plant growth conditions

Non-transgenic (NT) plantlets and two independent miR156-overexpressing lines of *P. edulis* (*Pe*OE-3 and *Pe*OE-5) and *P. cincinnata* (*Pc*OE-1 and *Pc*OE-4) were subcultured monthly on MS-based medium without growth regulators. For initial acclimatization, plants were transferred to 150 ml disposable cups containing horticultural soil conditioner substrate (Tropstrato^®^ HT Hortaliças, Vida Verde Indústria e Comércio de Insumos Orgânicos Ltda, Mogi Mirim, SP, Brazil), and kept in a growth room with a photoperiod of 16 h, irradiance of 50 μmol m^-2^ s^-1^ and temperature of 25 ± 1 °C for two weeks. Plants were subsequently transferred to 5-L polyethylene pots with Tropstrato^®^ HT, watered daily and maintained at an 11/13-h light/dark photoperiod and 25/16 °C (day/night) under greenhouse conditions. The access to genetic heritage adhered strictly to current Brazilian biodiversity legislation and was approved by the Brazilian National System for the Management of Genetic Heritage and Associated Traditional Knowledge (SISGEN) under permission number AF1B74A.

### Flow cytometry analysis

The genetic stability of miR156-OE lines was evaluated by assessing the relative DNA content from plants after 120 days grown in greenhouse. Briefly cell nuclei suspensions were prepared by cutting approximately 20-30 mg of young leaves from the samples and the internal reference (*Pisum sativum* ‘Citrad’, 2 C = 9.09 pg) and mixed with 1 mL of WPB isolation buffer (Doležel et al.,1998; Loureiro et al., 2007). The generated suspension was filtered through a nylon membrane (50 µm mesh) and 25 µL of propidium iodide solution (1 mg mL^-1^; Sigma-Aldrich, St. Louis, MO) and 5 mL RNAse (Amresco, Framingham, MA were added to each sample that were incubated in the dark for 1 h. The analysis was performed using a CytoFLEX flow cytometer (Beckman Coulter, CA). Three replicates were used, and fluorescence was quantified in at least 10,000 nuclei. Histograms were generated and analyzed using CytExpert 2.0 (Beckman-Coulter) and DNA content (pg) was calculated according to the formula proposed by Doležel and Bartoš (2005).

### Analyses of plant growth parameters

The main branch height (cm), internode length fourth to fifth leaf (shootwards) and branching index (ratio between the total length of lateral ramification and the length of the main plant axis; Morris et al., 2001) were assessed after 35 and 60 days of growth after transfer to greenhouse conditions for *P. edulis* and *P. cincinnata*, respectively.

### Number, size and ant visitation of extrafloral nectaries (EFNs)

The number of petiolar and laminar EFNs from fully expanded leaves was determined, considering fourth (4^th^) and sixteenth (16^th^) phytomer for *P. edulis*, and fourth (4^th^) twelfth (12^th^) phytomer for *P. cincinnata*. To determine the size, the nectaries were photographed in a stereomicroscope (Olympus SZX7-Olympus Corporation, Tokyo, Japan) and had the diameter (mm) and area (length x width-mm^2^) quantified using ImageJ software v. 1.43u (National Institutes of Health, USA) (Schneider et al., 2012). The number of ants visiting each plant was assessed at noon for ten consecutive days (Izaguirre et al., 2013). The number of ants on the plant daily was divided by the total number of branches greater than 1 cm. The experiments were repeated three times.

### Histological analyses and scanning electron microscopy (SEM)

Petiolar and laminar EFNs were fixed in Karnovsky (Karnovsky, 1965) from fully expanded leaves considering the sixteenth (16^th^) phytomer for *P. edulis*, and twelfth (12^th^) phytomer for *P. cincinnata*; and dehydrated in graded ethanol series (10-100%) and included in methacrylate (Historesin^®^, Leica Instruments, Germany). Sections (5 µm) were obtained using automatic feed rotary microtome (RM2155, Leica Instruments, Germany) and stained in toluidine blue (pH 4.7) for 2 min (O’Brien & McCully, 1981). The slides were mounted in Permount^®^ SP15-500 (Fisher Scientific, Waltham) and observed under a light microscope (AX70 TRF, Olympus Optical) with U-photo system, coupled to a digital camera (Spot Insightcolour 3.2.0, Diagnostic Instruments Inc.). For SEM, samples were fixed in Karnovsky, submitted dehydration in ethylic series, and dried at the critical point, utilizing CO2 in Balzers equipment (model CPD 020, Bal-Tec, Balzers, Liechtenstein). Subsequently, the fragments were covered in gold using the cathodic spraying process in Sputter Coater equipment (Model FDU 010, Bal-Tec, Balzers, Liechtenstein) (Silveira,1989). The observations and photographic documentation were carried out using a scanning electron microscope (model Leo 1430 VP, Zeiss, Cambridge, England). The number of cells constituting the nectary epidermis and parenchyma (nectariferous and subnectariferous parenchyma) was determined using Image J software (Schneider et al., 2012).

### Total RNA extraction

To analyze the expression dynamics of miR156 in consecutive phytomers in *P. cincinnata*, fully expanded leaves were collected, according to Silva et al. (2019). For RNA-seq and expression analysis of selected genes and microRNAs, young leaf primordia from the 16^th^ phytomer for *P. edulis* and the 12^th^ phytomer for *P. cincinnata* were used. Total RNA was extracted from 30 mg of macerated leaf tissue using Trizol according to the manufacturer’s instructions (TRIzol™ Reagent – Invitrogen). RNA quantification was performed with the aid of NanoDrop^TM^ Lite (Thermo Scientific, Wilmington, DE) under an absorbance of 260 nm, and the integrity of the RNA samples was verified by 1.5% agarose gel electrophoresis (RNAse free). Samples were then treated with DNAse (RQ1 RNAse-Free DNAse-Promega), according to the manufacturer’s instructions.

### Library Preparations and RNA sequencing

Total RNA and Dynabeads^®^ Oligo (dT) 25 (Thermo Fisher Scientific, Waltham, MA) were used to isolate mRNA. The resulting mRNA fragments of ∼400 nucleotides were converted to double-stranded complementary DNA (cDNA) using random hexamer primers and corresponding enzymes following standard DNA library protocols for sequencing. Briefly, cDNA was end-repaired, phosphorylated, and adenylated. Common TruSeq adapters containing 8-bp indexes (i5 and i7) suitable for Illumina sequencing were then ligated to the adenylated molecules, and the resulting libraries were amplified by 13 cycles of PCR to enrich for properly ligated molecules. The final libraries were quantified using PicoGreen (Thermo Fisher Scientific) and equally combined into a single sample, then sequenced on an Illumina HiSeq X (Illumina Inc., San Diego, CA) instrument. Paired-end reads with an average length of 150 bp were obtained. Library preparation and sequencing were carried out by Rapid Genomics, LLC (Gainesville, FL).

### *De novo* assembly and annotation

For each genotype, three and four RNA libraries were sequenced for *P. edulis* and *P. cincinnata*, respectively. Trimmomatic software v 0.39 removed adapters and reads with phred score quality less than Q20. All cleaned reads from each pair-end were pooled and assembled using Trinity software version 2.11.0 with default parameters. The coding DNA sequence (CDS) was predicted using transdecoder v5.5.0 software (https://github.com/TransDecoder/TransDecoder) with default parameters. For the transcriptome assessment, we used BUSCO (Benchmarking Universal Single-Copy Orthologs) (Manni et al., 2021).

All assembled transcripts and predicted ORFs were annotated against Swiss-prot, Uniref 50, Uniref90, Pfam, EggNOG, Gene Ontology (GO) databases using trinotate pipeline. For BLASTp search, we used a cut-off e-value of 1.0E-3. All results were deposited into SQLite database, and a spreadsheet summary report was generated. To quantify the gene expression level of all samples, we performed mapping clean reads to assembled transcripts using the salmon v 0.14.1 program. From this abundance estimation, we used the DESeq2 v1.36.0 R package to carry out the differential expression analysis of gene and transcript level of trinity software. Heatmap and volcano were plotted in R v 3.5.1. GO enrichment analysis of a subset of DEGs (*p < 0.001*) was conducted using Goseq, GO.db, and qvalue packages from R with default parameters.

### Identification, phylogenetic and sequence analysis of *Passiflora SPL* gene family

To identify the sequences containing the SBP/SPL domain (PF03110), all 58087 transcripts generated by RNA from *P. edulis* and 71034 from *P. cincinnata* annotated against the Swiss-prot database were used. Transdecorder predicted a total of 112 sequences of *SPL* genes for *P. edulis* and 90 sequences for *P. cincinnata*, which were used as a query to conduct BLASTx in the NCBI (National Center for Biotechnology Information) database. 11 *SPL* family genes in *P. edulis* and 11 in *P. cincinnata* were selected considering the high identity and coverage of the sequences. All retrieved CDS were translated in all six frames using EMBOSS Transeq (https://www.ebi.ac.uk/Tools/st/emboss_transeq/) and analyzed by PFAM v35.0 (Salazar et al., 2020) to maintain only those containing the SBP/SPL domain (Supplementary Table S4). All unique and primary coding sequences from *Solanum lycopersicum* (ITAG4.0 assembly), *Populus trichocarpa* (v4.1 assembly), *Vitis vinifera* (v2.1 assembly), and *Panicum virgatum* (v4.1 assembly) were downloaded at Phytozome v13 and used to recovery sequences containing PF03110 domain. The name of SBP or SPL gene names were used as described previously for *Arabidopsis* (Preston & Hileman, 2013), tomato (Salinas et al., 2012), poplar (Guo et al., 2021; Hou et al., 2013) grapes (Hou et al., 2013; Díaz-Riquelme et al., 2012) and switchgrass (Wu et al., 2016).

All protein sequences for each multigene family were aligned using the default settings of Multiple Sequence Comparison by Log-Expectation (MUSCLE) (http://www.ebi.ac.uk/Tools/msa/muscle/; Madeira et al., 2022). A Maximum Likelihood (ML) phylogenetic analysis was performed as using PhyML3.0 (Guindon et al., 2010), using Smart Model Selection (SMS) to select the best model (Lefort et al., 2017) (like VT+R+F substitution model) and aLRT branch support testing (Anisimova & Gascuel, 2006). The phylogenetic tree was visualized using the iTOL software (https://itol.embl.de; Letunic & Bork, 2021).

High-confidence prediction of miRNA156 targets in *P. edulis* was performed by *psRNATarget* (Dai et al., 2018), using the database https://www.zhaolab.org/psRNATarget/analysis. We have used a strict cut-off threshold for the Maximum Expectation (E) parameter (range 0 - 3.0).

### Analysis of gene expression by RT-qPCR

For microRNA quantification, we used the stem-loop qPCR protocol previously described by Varkonyi-Gasic et al. (2007). For the remaining genes, cDNA was synthesized using the SuperScript III First-Strand Synthesis System for the RT-PCR kit (Invitrogen, San Diego, CA), according to the manufacturer’s instructions. P*C*R reactions were performed using iTaq Universal SYBR Green supermix (Bio-Rad) and analyzed in a Step-OnePlus real-time PCR system (Applied Biosystems). The RT-qPCR analyses used two or three biological samples with two technical replicates each. Expression levels were calculated relative to the housekeeping gene *PeACTIN1* (Cutri & Dornelas, 2012) using the 2^-ΔΔCt^ method (Livak and Schmittgen, 2001). The primers used were designed to a unique sequence within the predicted coding region for each gene from sequences obtained by RNA-seq of *P. edulis* and *P. cincinnata* and are listed in Supplementary Table S1.

### Analysis 5’ Rapid amplification of cDNA ends (RACE)

Five micrograms of total RNA from leaves of *Passiflora edulis* was treat with DNase I (Invitrogen) and ligated to an RNA adapter following the manufacture’s guide of GeneRacer kit (Invitrogen), the RNA ligation reaction was performed for 5 hours at 37 °C. The complete amount of RNA was reverse transcribed to cDNA using the oligodT from the GeneRacer kit and following the manufacture’s guide of ImProm-II Reverse Transcriptase (Promega). The *PeSPL6* and *PeSPL13a* 5’-ends were amplified by PCR using the GoTaq Master mix (Promega), the forward 5’ Primer from GeneRacer kit and the reverse Gene Specific Primer (GSP). The PCR product was used as a NESTED PCR template using the forward 5’ Nested Primer from GeneRacer kit and the reverse Nested Gene Specific Primer (Nested GSP). The fragments size were confirmed by agarose gel (1%), the PCR fragments was purified using the Qiaquick PCR purification kit (Qiagen) and cloned into pGEM-T easy (Promega) vector. Nine independent clones from *PeSPL6* and two from *PeSPL13a* were sequenced using the forward M13 primer.

### Sugar quantification

A pool of petiolar and laminar EFNs tissues isolated from fully expanded leaves was collected at midday. The samples were then macerated in liquid nitrogen and subjected to five successive treatment steps with 80% ethanol at 100°C for 5 min. After each extraction, the mixture was centrifuged (Eppendorf 5415D centrifuge) at 14000 rpm for 5 min. The total ethanolic extract obtained was evaporated at 50 °C and the sugars resuspended in 1 mL of 80% alcohol, subsequently the solution was filtered through an Econofilter membrane filter (Agilent Technologies 0.2 μm). The filtrate was analyzed by high performance liquid chromatography (HPLC) (Shimadzu^®^ series 20A, Kyoto, Japan), equipped with a column (Shim-pack GIST NH2 Shimadzu) and refractive index detector, containing the aminopropyl group (-NH2) as stationary phase. A mixture of acetonitrile and water (75:25) under isocratic conditions was used as a mobile phase. The analyses were performed at 40°C, under flow rate of 1 mL min^-1^. Sucrose, fructose and glucose were quantified using a calibration curve that correlates peak areas and known concentrations of the analyte.

### Statistical analysis

The experimental design was completely randomized. The analysis of variance (*P < 0.05*) was used to determine the effects of the treatments: for the number and size of extrafloral nectaries, the means were submitted to the Tukey test (*P ≤ 0.05*), and for the other variables, the Student’s t-test (*P ≤ 0.05*) was applied using the program R-Bio^®^ version 171 (hptts://biometria.ufv.br/) (Bhering, 2017). Graphs were generated in Rstudio^©^ v.1.4 software using the ggplot2 package.

## Results

### High miR156 levels dramatically alters morphometric parameters and leaf morphology in *Passiflora* species

The age-dependent morphological changes observed in *P. edulis* leaves are associated with a temporal decline in miR156 expression and a concomitant increase in *PeSPL9* transcript levels (Silva et al., 2019). Similarly, in the early stages of *P. cincinnata* vegetative development, young leaves were trilobed and had elevated levels of miR156 (Fig. S1a, d). As vegetative development progressed, leaves gradually became five-lobed and the levels of miR156 were strongly reduced. In constrast, *PcSPL9* and miR172 transcript levels increased (Fig. S1 b-d), similarly to what we previously observed in *P. edulis* (Silva et al., 2019). In both in *P. edulis* and *P. cincinnata*, the presence of laminar EFNs was correlated with reduced miR156 levels and a subsequent increase in *SPL* transcript levels (Silva et al., 2019; Fig S1c), which reinforces the hypothesis that the miR156-*SPL* module may have been co-opted to regulate EFN emergence, independent of the leaf shape.

To comprehensively investigate the role of miR156-targeted *SPLs* in leaf heteroblasty and EFN formation in *Passiflora* species, we generated independent transgenic events overexpressing miR156 (hereafter referred to as miR156-OE or OE) in *P. edulis* (*Pe*OE) and *P. cincinnata* (*Pc*OE; Fig. S2a). Nuclear DNA content was maintained in all miR156-OE lines relative to non-transgenic or control plants (NT: *Pe*NT and *Pc*NT) (Fig. S2 b-d), confirming no modifications in transgenic *Passiflora* ploidy. *P. edulis* and *P. cincinnata* miR156-OE plants showed altered vegetative architecture and exhibited a moderate bushy appearance (Fig. 1a). The miR156-OE plants in both species exhibited reduced height and shorter 4^th^ phytomer internode length than NT (Fig. 1b-c). Additionally, we observed increased miR156-OE axillary shoot growth, leading to increased branching index in transgenic *P. edulis* (∼3-fold) and *P. cincinnata* (∼1.5-fold) when compared with NT (Fig. 1d).

**Figure 1.**
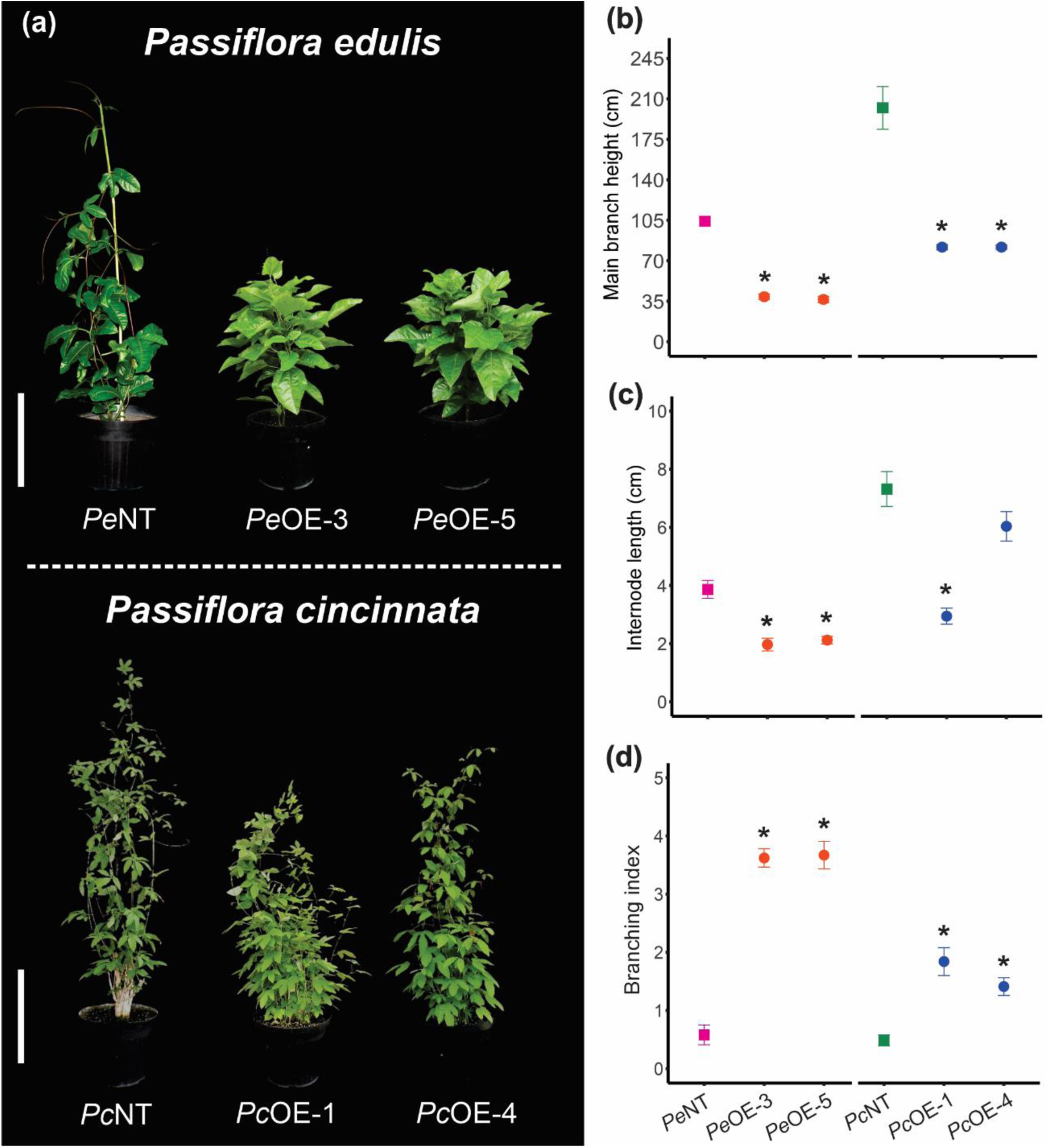
Shoot architecture is modified in miR156-overexpressing *Passiflora* species. **(a)** Vegetative architecture of non-transformed (*Pe*NT and *Pc*NT) and plants overexpressing miR156 (miR156-OE) in two distinct lines of *Passiflora edulis* (*Pe*OE-3 and *Pe*OE-5) on the 35^th^-day post acclimatization (DPA) and *Passiflora cincinnata* (*Pc*OE-1 and *Pc*OE-4) on the 60^th^ DPA in a greenhouse. Scale Bar: 30 cm. (n=5). **(b)** Main stem height. **(c)** Internode length at 4 ^th^ node. **(d)** Branching index. Values are presented as mean ± SE (n=8). \**P* ≤ 0.05 (Student’s *t*-test).

Seed-derived *P. edulis* plants exhibit monolobed leaves during the juvenile phase, and, at the 10^th^ node, these leaves become trilobed (Silva et al., 2019). On the other hand, *in vitro-*derived *Pe*NT plants exhibited leaf trilobing at the 6^th^ node. In contrast, *Pe*OE plants produced consistently monolobed up to the 10^th^ node and only slightly lobed leaves from the 11^th^ node (Fig. 2a). In *P. cincinnata* plants derived from *in vitro* explants, five-lobed leaves with completely distinct lobes occurred between the 6^th^ to 7^th^ nodes. Conversely, five-lobed leaf production was delayed in *Pc*OE plants (7^th^ to 8^th^ nodes) and lobing was consistently shallower and less distinct than in *Pc*NT plants (Fig. 2b). These results indicate that the role of the miR156-*SPL* module in regulating leaf heteroblasty is conserved in *Passiflora* species.

**Figure 2.**
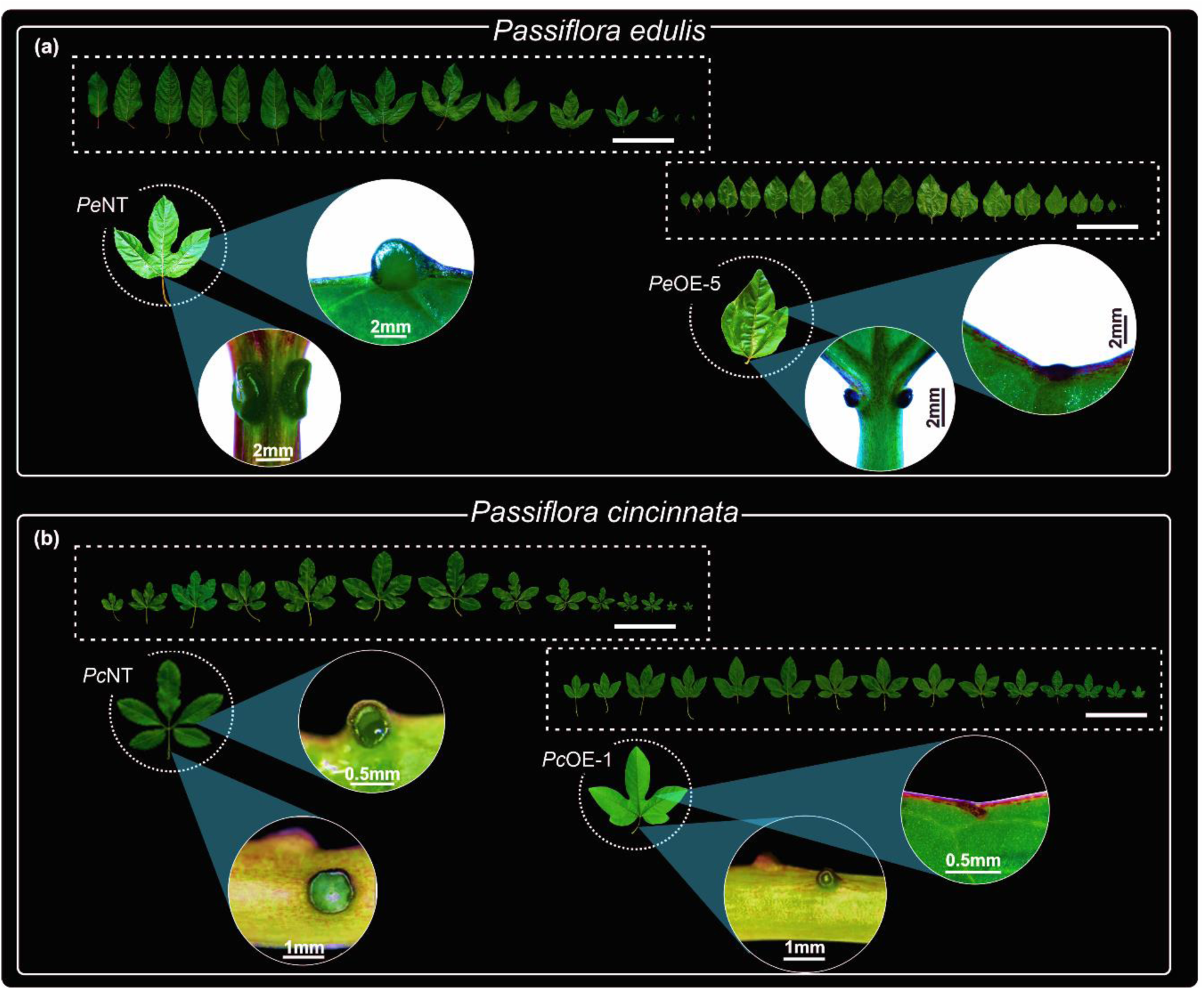
High levels of miR156 alter leaf heteroblastic development and extrafloral nectary (EFN) formation in *Passiflora* species. (**a**) Leaf morphology and EFN distribution in the petioles and leaf lamina of non-transformed (*Pe*NT) and over-expressing miR156 (*Pe*OE-5 plants of *Passiflora edulis* and **(b)** non-transformed (*Pc*NT) and over-expressing miR156 (*Pc*OE-1) plants of *Passiflora cincinnata*. Details of EFN of leaf 15 of *Pe*OE-5 line and leaf 12 of *Pc*OE-1. Scale Bar: 15 cm.

### Extrafloral nectary size and abundance are age-dependent in *Passiflora* species

To determine whether miR156 levels affect EFN formation, we carried out a detailed phenotypic analysis of EFN development in *P. edulis* and *P. cincinnata* plants constitutively expressing miR156. While *Passiflora* petiolar EFNs are paired and located on the adaxial side of the petiole, laminar EFNs are located abaxially on the leaf margin in the sinuses between leaf lobes (Fig. S1d; Fig. 2). For both species, miR156-OE EFNs were underdeveloped or even absent in some cases when compared with NT plants (Fig. 2).

Reduced miR156 expression is correlated with the appearance of EFNs in some *Vachellia* species (Leichty & Poethig, 2019). To examine the relationship between shoot identity and the formation of petiolar and laminar EFNs in *Passiflora* species, we performed a detailed analysis of these structures in different vegetative phases of NT (control) and miR156-OE plants. *Pe*NT plants exhibited a single pair of petiolar EFNs on both the 4^th^ (juvenile) and the 16^th^ (adult) leaves; however, these EFNs were significantly larger on adult leaves (Fig. 3 a-c). *Pe*OE plants also had two petiolar EFNs on nodes 4 and 16, but they were smaller than *Pe*NT nectaries on the same nodes and did not increase in size during vegetative development. Additionally, while *Pe*NT plants had four laminar EFNs on the 16^th^ leaf (Fig. 3d), *Pe*OE plants had only one relatively small EFN (Fig. 3 d-f).

**Figure 3.**
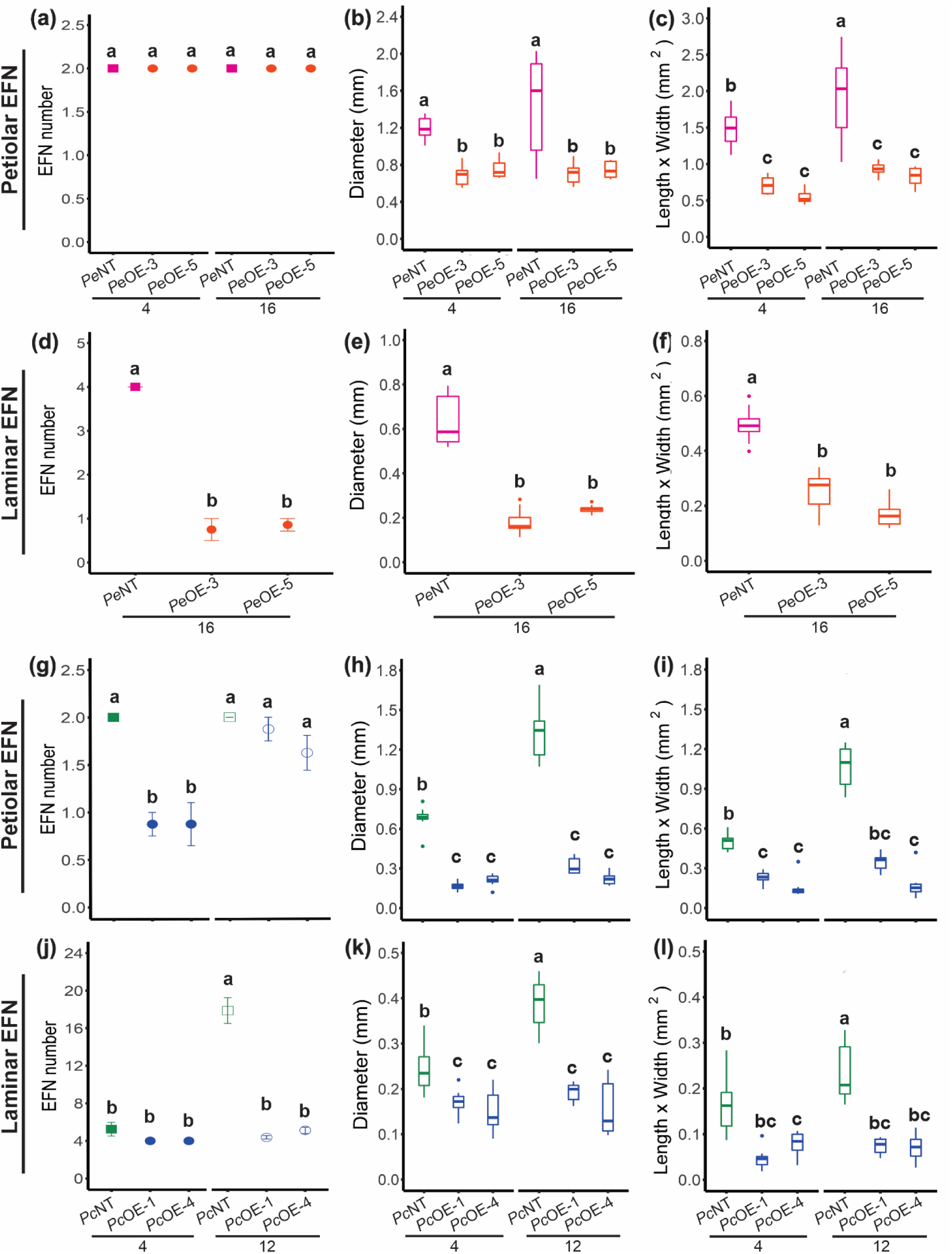
Extrafloral nectary (EFN) size and number are age-dependent in *Passiflora* species. **(a)** Number of petiolar EFN in non-transformed (*Pe*NT) and miR156-overexpression (*Pe*OE-3 and *Pe*OE-5) lines of *Passiflora edulis.* **(b-c)** Diameter and area (Length x Width) of petiolar EFN. **(d)** Number of laminar EFN in *Pe*NT, *Pe*OE-3, and *Pe*OE-5 plants. **(e-f)** Diameter and area of laminar EFN. **(g)** Number of petiolar EFN in non-transformed (*Pc*NT) and *Passiflora cincinnata* miR156-overexpression (*Pc*OE-1 and *Pc*OE-4) lines. **(h-i)** Diameter and area of petiolar EFN. **(j)** Number of laminar EFN in *Pc*NT, *Pc*OE-1 and *Pe*OE-4 plants. **(k-l)** Diameter and area of laminar EFN. Numbers 4 and 16 in *P. edulis* and 4 and 12 in *P. cincinnata* correspond to leaf position (node). Values are presented as mean ± SE (n=8). \**P* ≤ 0.05 (using ANOVA followed by Tukey’s pairwise multiple comparisons).

In *P. cincinnata*, petiolar and laminar EFNs were found at both juvenile and adult vegetative stages. *Pc*NT plants had paired petiolar EFNs on the 4^th^ (juvenile) and 12^th^ (adult) leaves, but the nectaries were larger on leaf 12 (Fig. 3 g-i). *Pc*OE plants had fewer petiolar EFNs on leaf 4, but there was no difference in petiolar EFN number on leaf 12 compared with *Pc*NT (Fig. 3g). The size of the petiolar EFNs on leaves 4 and 12 of *Pc*OE plants was 4- and 6-fold smaller than the nectaries of *Pc*NT plants, respectively (Fig. 3h-i). Both *Pc*NT and *Pc*OE leaves had an average of five laminar EFNs on leaf 4 (Fig. 3j), but they were smaller in diameter in *Pc*OE than *Pc*NT (Fig. 3 k-l). In contrast, *Pc*NT plants had an average of 18 laminar EFNs on leaf 12, whereas *Pc*OE plants had only 4-5 EFNs (Fig 3 j-l). *Pc*OE laminar EFNs were also consistently smaller than *Pc*NT nectaries and did not increase in size. In summary, constitutive expression of miR156 led to an overall decrease in the number and size of *Passiflora* EFNs. Collectively, these observations suggest that the initiation and growth of EFNs are age-dependent in *Passiflora* species.

To determine how miR156 affects EFN growth during early development, we next examined in detail the morphoanatomy of EFNs at the cellular level from fully expanded adult leaves in NT and miR156-OE plants. In *Pe*NT, the laminar EFN has an elliptical shape with a concave central region (Fig. 4a; Lemos et al., 2017), whereas in *Pe*OE plants, the laminar EFNs developed as a protuberance with a convex surface (Fig. 4b-c). Although all three characteristic nectary tissues (nectary epidermis, nectariferous parenchyma, and subnectariferous parenchyma; Cardoso-Gustavson et al., 2013; Macêdo et al., 2021; Moraes et al., 2022) were observed in *Pe*NT and *Pe*OE plants (Fig. 4d-f), the number of cells in each layer was strongly reduced in these plants (Fig. 4m). In *P. cincinnata*, laminar EFNs have an obconical shape, curved towards the abaxial region of the leaves (Fig. 4g). In contrast, *Pc*OE-1 and *Pc*OE-4 exhibited an elongated cylindrical shape that featured an apex with a convex surface or a small central cavity (Fig. 4 h-i). Similar to *P. edulis*, tissue patterning was comparable between *Pc*NT and *Pc*OE (Fig. 4 j-l), but *Pc*OE laminar EFNs exhibited reduced cell numbers in each tissue type (Fig. 4n).

**Figure 4.**
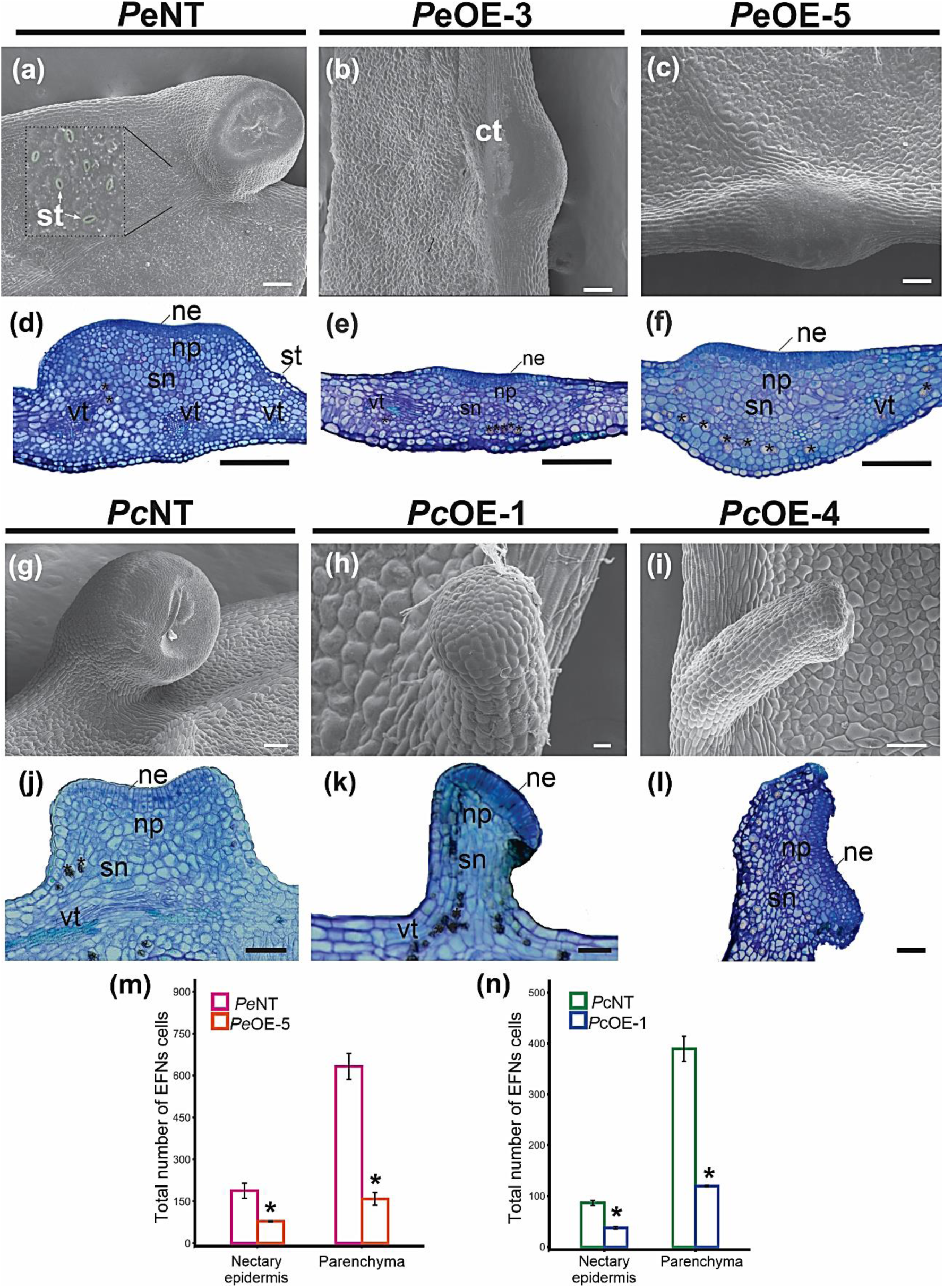
Cell number, but not patterning, of laminar extrafloral nectaries (EFN), is affected by high levels of miR156 in *Passiflora* leaves. Scanning electron microscopy images in *Passiflora edulis*: **(a)** Laminar EFN nectary in non-transgenic (*Pe*NT) plants and **(b)** miR156-overexpression *Pe*OE-3 and (c) *Pe*OE-5 plants. Note the difference in morphology between non-transgenic and miR156-OE plants. ct: cuticle, st: stomata. Light microscopy in *P. edulis* plants: **(d)** Transverse section of the leaf showing a laminar EFN in *Pe*NT and (e) *Pe*OE-3 and (f) *Pe*OE-5 with three distinct regions: Nectary epidermis (ne), nectariferous parenchyma (np); subnectariferous parenchyma (sn); leaf vascular tissue (vt) and cells containing calcium oxalate crystals (**\***). Scanning electron microscopy images in *P. cincinnata*: **(g)** Laminar EFN in non-transgenic (*Pc*NT) plant showing their secretory surface located at a concavity at medium-top region. **(h)** Laminar EFN showing an elongated shape in overexpressing miR156 plants *Pc*OE-1 and (i) *Pc*OE-4. Light microscopy **(j)** Transverse section of the leaf showing a laminar EFN in *Pc*NT and (k) *Pc*OE-1 and (l) *Pc*OE-4 plants showing the three distinct tissues. **(m)** Total number of cells of nectary epidermis and parenchyma (necteriferous + subnectariferous) of the laminar EFNs in *P. edulis* and (n) *P. cincinnata*. Values are presented as mean ± SE (n=3). \**P* ≤ 0.05 (Student’s *t*-test). Scale Bars: a-c =100 μm, d-f = 50 μm, g=100 μm, h = 20 μm, i-l= 100 μm.

Petiolar EFN in *P. edulis* displays an elliptical shape with a small cavity in the center (Fig. S3a), through which nectar is released (Lemos et al., 2017). *P. edulis* miR156-OE plants showed petiolar EFNs with an elliptical shape similar to *Pe*NT, except for the cavity in the center of the nectary (Fig. S3 b-c). Small elevations in the central region of the EFN were observed in *Pe*NT and *Pe*OE plants, probably representing nectar secretion that was retained in the subcuticular space (Fig.S3 a-b). Petiolar EFNs in *Pc*NT plants also displayed an elliptical shape with a central concavity from which nectar is exuded (Fig. S3j). In *Pc*OE plants, on the other hand, petiolar EFNs exhibited an elongated cylindrical -shape with a small central concavity (Fig. S3 k-l). Similar to laminar EFNs, petiolar EFNs of NT and miR156-OE *Passiflora* plants are composed by nectary epidermis, nectariferous and subnectariferous parenchyma (Fig. S3 d-i; m-s). Although NT and miR156-OE plants in *P. edulis* and *P. cincinnata* showed similar tissue patterning, the morphology and number of cells constituting the secretory tissue of petiolar and laminar EFNs were likely reduced in the miR156-OE plants, suggesting that cell division is compromised in miR156-overexpressing leaves.

### Transcriptional reprogramming of miR156-overexpressing *Passiflora* leaf primordia

To better understand the molecular mechanisms by which miR156 modulates leaf and EFN development, we employed RNA-seq to monitor changes in gene expression in *P. edulis* and *P. cincinnata* NT and miR156-OE leaf primordia, prior to the appearance of petiolar and laminar EFNs. After filtering low-quality reads, a total of 212,141 transcripts with average 677.33 bp and N50 equal to 1628 bp for *P. edulis* and 239,343 transcripts with average 629.10 bp and N50 equal to 1444 bp for *P. cincinnata* were assembled. Our functional annotation indicated that 58,087 ORFs (76.3%) and 63,376 ORFs (76.9%) were annotated in the SWISS-PROT database for *P. edulis* and *P. cincinnata*, respectively. The RNA-seq analysis detected 1048 differentially expressed genes (DEGs) in *P. edulis* (FDR < 0.05 and fold change >1.5), including 407 up-regulated and 641 down-regulated genes (Fig. S4a; Table S2). In *P. cincinnata*, a total of 667 DEGs were identified, including 316 up-regulated and 351 down-regulated genes (Fig. S4b; Table S3). Gene Ontology (GO) analysis classified DEGs (P < 0.05) into three categories, biological process, cellular component and molecular function (Fig. S5). In general, DEGs were more enriched in the biological process category, which included genes associated with flower development (GO: 0009908), regulation of vegetative phase change (GO: 0010321), development of the shoot reproductive system (GO: 0090567) and senescence of floral organs (GO: 0080187).

To characterize the effects of constitutive miR156 expression on *SPL* transcript levels we first interrogated our RNA-seq data to identify *SPL* sequences using TransDecoder (https://github.com/TransDecoder/TransDecoder). This revealed a set of eight *P. cincinnata* and nine *P. edulis SPL* members, referred to as *PcSPLs* and *PeSPLs* respectively, detected across our samples (Fig. S6a). Phylogenetic analyses demonstrated that these genes represented all the major clades of *SPL* family members previously described. *PeSPL/PcSPL* genes were annotated based on the closest phylogenetic relationship with *Arabidopsis* and grape *SPLs* (Salinas et al., 2012; Preston & Hileman, 2013; Table S4). Five *SPLs* (*PcSPL6*/*PeSPL6*, *PcSPL9*, *PcSPL13a*/*PeSPL13a*, *PcSPL13b*/*PeSPL13b*, *PeSPL13s*) contain the target site for miR156 in their transcript sequences (Fig. S6b). We did not find the miR156 recognition site for all orthologs of *SPL* genes targeted by miR156 in other angiosperms due to incomplete transcript sequences in our dataset (Fig. S6a, c; Table S4). Among the identified miR156-targeted *SPLs*, *SPL6, -9,* and *-13a* were strongly repressed in miR156-OE lines relative to NT plants in both species (Fig. 5a-f), suggesting they are strong candidate regulators of heteroblastic leaf development in *Passiflora*. To determine whether miR156 regulates any non-*SPL* targets we employed psRNATarget (Dai et al., 2018). Similar to other species we found no well-supported targets outside of the *SPL* family (Table S4). To confirm that miR156 directly targets *Passiflora SPL* transcripts we identified miR156 cleavage sites in *PeSPL6* and *PeSPL13a* by 5’ RACE (Fig. 5g). This would suggest that, similar to *Arabidopsis* (He et al., 2018), miR156 functions at least in part through transcriptional cleavage in *Passiflora*.

**Figure 5.**
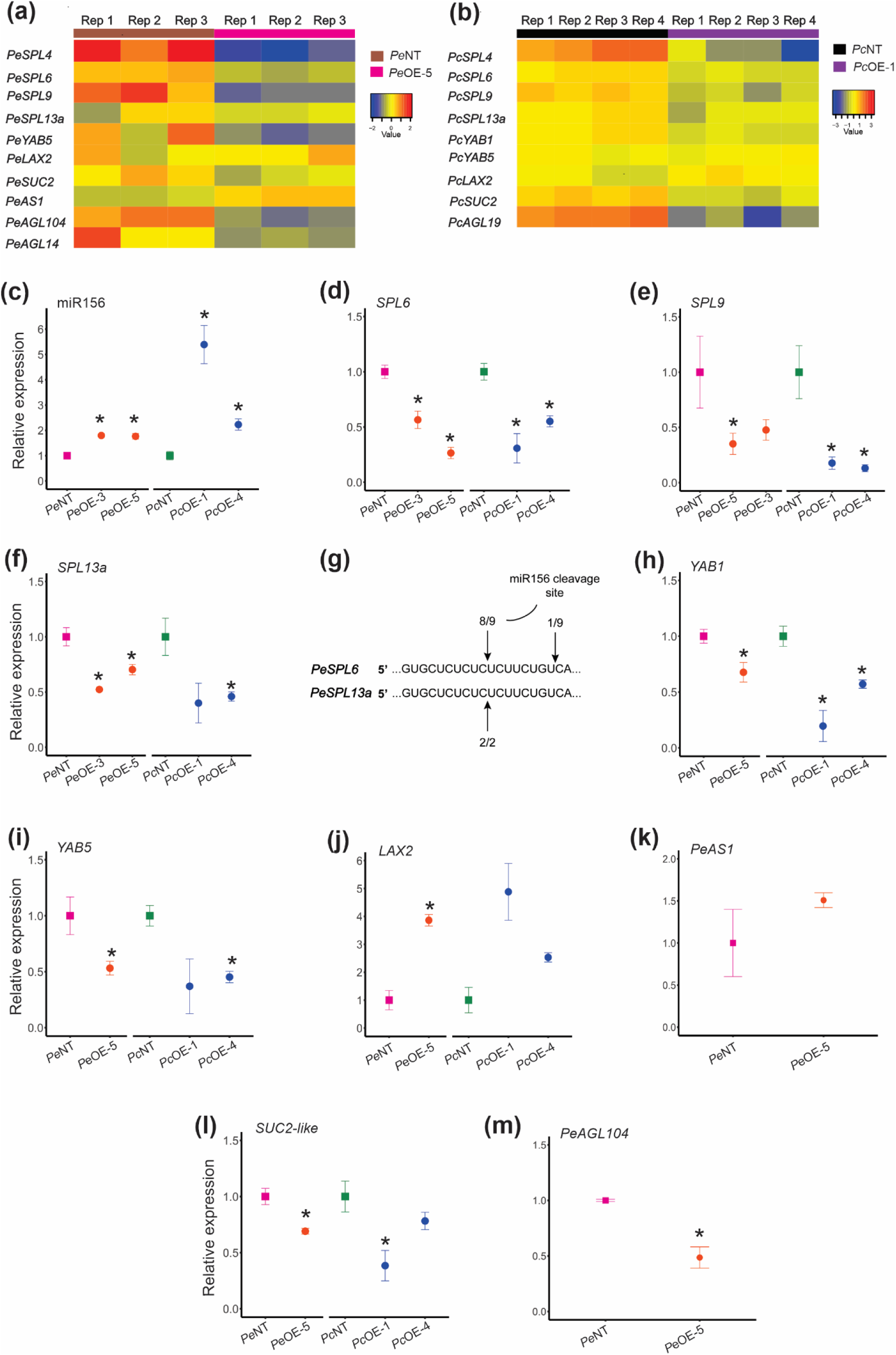
miR156 overexpression modulates the levels of *SPLs* and genes associated with *Passiflora* leaf ontogeny and nectary development. (**a, b**) Identification of differentially expressed genes (DEGs) up-regulated (red) and down-regulated (blue) by RNA-seq analysis in leaf primordia of non-transgenic (NT) plants compared with overexpressing miR156 (miRNA-OE) plants in *Passiflora edulis* (a) and *P. cincinnata* (b). R1 to R4 indicate biological replicates. **(c)** Relative expression by RT-qPCR analysis of miR156 in *P. edulis* (*Pe*NT, *Pe*OE-3, *Pe*OE-5) and *P. cincinnata* (*Pc*NT, *Pc*OE-1, *Pc*OE-4), *SPL6* (d), *SPL9* (e), and *SPL13a* (f). (g) Modified 5’-RACE analyses of cleaved *PeSPL6* and *PeSPL13a* transcripts in leaf tissue. The 5’-ends of *PeSPL6* and *PeSPL13a* map to the miR156 recognition site. **(h)** Relative expression of *YAB1*, *YAB5* (i), *LAX2* (j), *AS1* (k), *SUC2-like* (l), and *AGL104* (m). Values are mean ± SE (n=3). \**P* ≤ 0.05 (Student’s *t*-test).

Overexpression of miR156 and the consequent reduction in *SPL* activity in *Passiflora* impacted the expression of a number of genes associated with leaf ontogeny, such as *YABBY1* (*YAB1*) and *YAB5*. Members of the YABBY gene family, including the floral nectary-enriched *YAB5* and CRABS *CLAW* (*CRC*), are expressed abaxially in leaves and floral organs and are responsible for establishing or interpreting organ polarity in *Arabidopsis* (Siegfried et al., 1999; Bowman & Smyth, 1999; Eshed et al.,1999). While we did not detect *CRC* transcripts in our samples of young leaves, *YAB1* and *YAB5* were downregulated in miR156-OE leaf primordia in both species (Fig. 5 a, b, h, i; Table S2, S3). On the other hand, orthologs of *AUXIN TRANSPORTER-LIKE PROTEIN 2* (*LAX2*), which regulates leaf margin serration in *Arabidopsis* (Kasprzewska et al., 2015) and *ASYMMETRIC LEAVES1* (*AS1*), which regulates dorsoventral polarity (Husbands et al., 2015), were upregulated in *P. edulis* miR156-OE lines (Fig. 5j, k; Table S2, S3). Interestingly, the *SUCROSE TRANSPORT PROTEIN 2* (*SUC2*) gene, responsible for sucrose transport (Chandran et al., 2003), was downregulated in miR156-OE plants (Fig. 5l). In other species members of the *AGAMOUS-like MADS-box* (*AGL*) family of genes have been shown to promote leaf complexity and floral nectary development (Kram et al., 2009; Ostria-Gallardo et al., 2016; Chen et al., 2021). Consistent with these observations, *Passiflora AGL14*, *AGL19* and *AGL104* were all downregulated in miR156-OE leaf primordia (Fig. 5a, b, m), which suggests that *SPLs* may be required for their activation. Indeed, *AGLs* have been shown to be direct targets of miR156-targeted SPLs in *Arabidopsis* (Wang et al., 2009; Gao et al., 2018). Other genes necessary for floral nectary development (Chen et al., 2021), such as *LEAFY*, were also downregulated in *PeOE* leaf primordia (Table S2). Collectively, our data are consistent with the hypothesis that the miR156-*SPL* module has been coopted into the regulation of genetic networks associated with nectary development.

### Ecological relationships and sugar profile of extrafloral nectaries are affected in miR156-overexpressing *Passiflora*

*Passiflora* EFNs offer nectar to territorial and aggressive ants, establishing important mutualistic and ecologically impactful relationships (Cardoso-Gustavson et al., 2013). To determine whether *SPL* activity is important for the function of EFNs we evaluated ant visitation of both *P. edulis* and *P. cincinnata* plants in the greenhouse for 10 days at the period of highest ant activity. Visiting ants belonged to the genus *Brachymyrmex*, which are ∼3mm in lenght with an acidopore and 9-segmented antennae that lacks an antennal club (Fig. 6a; Ortiz-Sepulveda et al., 2019). Ant visitation was significantly reduced in *P. edulis* and *P. cincinnata* miR156-OE plants compared to *Pe*NT and *Pc*NT. (Fig. 6b-c).

**Figure 6.**
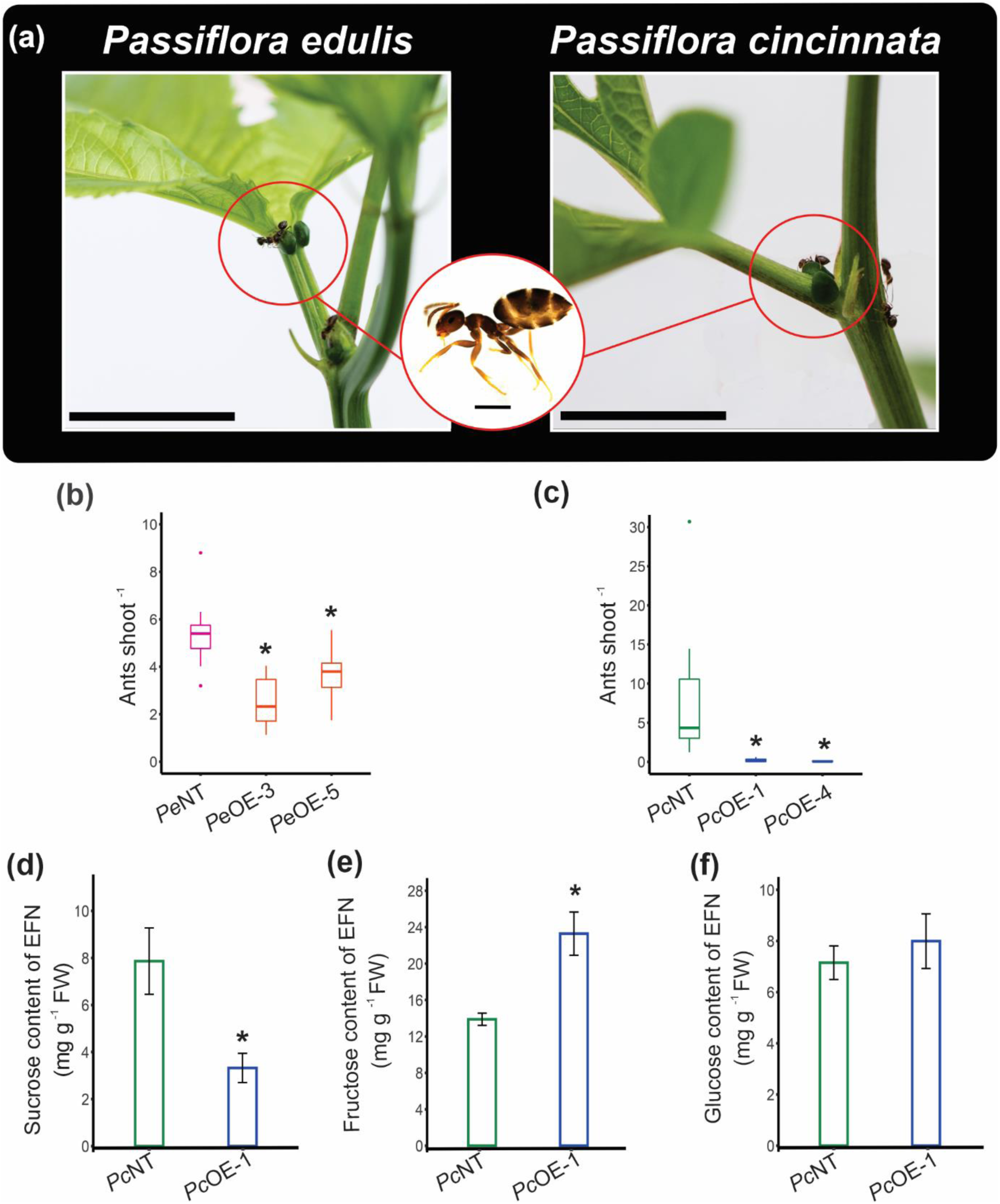
Reduced *SPL* expression modified sugar profiles in miR156-overexpressing *Passiflora* extrafloral nectaries (EFNs), and negatively impacted ant visitation. (**a**) Ants collecting nectar on petiolar EFNs of *Passiflora edulis* and *P. cincinnata*. Scale Bar:1cm. Detail of the ant *Brachymyrmex* spp. Scale Bar: 0.5 mm. **(b, c)** Ant visitation values per plant are presented as mean ± SE in non-transgenic (NT) and *P. edulis* overexpressing miR156 (*Pe*OE**-**3 and *Pe*OE**-**5) and (b) *P. cincinnata* (*Pc*OE**-**1 and *Pc*OE**-**4**) (c)** (n= 8-9). **(d)** Sucrose, **(e)** fructose and **(f)** glucose content of the laminar and petiolar EFN pool of *Pc*NT and *Pc*OE-1 (n = 3). \**P* ≤ 0.05 (Student’s *t*-test).

The nectar exuded by EFNs is rich in carbohydrates and is an attractant for food-seeking ants (Cardoso-Gustavson et al., 2013). EFNs in miR156-OE plants are smaller than in NT plants (Fig. 4), which might make them less attractive to ants. In addition, the quality of the nectar is impacted by the size of EFNs in different species, and affects ant visitation (Alencar et al., 2023). We therefore evaluated nectar availability and carbohydrate content in the EFNs from miR156-OE and NT plants. Although nectar availability was consistently observed in miR156-OE plants in both species during the evaluated period, the sucrose content of *Pc*OE EFNs was significantly lower than *Pc*NT (Fig. 6 d). On the other hand, fructose content was higher in *Pc*OE and glucose level was comparable with *Pc*NT nectaries (Fig. 6 e-f). In *P. edulis* EFNs, sucrose was not detected, and fructose and glucose content showed no significant differences between *Pe*OE and *Pe*NT, although *Pe*OE nectaries showed slightly higher fructose and lower glucose content compared to *Pe*NT (Fig. S7a,b). Collectively, our results indicate that low abundance and defective EFN patterning combined with modifications in carbohydrate composition of the nectar correlate with the reduced ant visitation observed in miR156-OE *Passiflora* plants. Thus, age-dependent miR156-targeted *SPLs* might be required for the ecological interaction potential of *Passiflora* species.

## Discussion

Temporal patterning of shoot development in *Passiflora* species likely requires the activity of miR156-targeted SPLs

Leaf heteroblasty is used as a taxonomic trait for several *Passiflora* species, and can distinguish between juvenile and adult phases of vegetative growth (Chitwood & Otoni 2017 a, b; Silva et al. 2019). Similar to *P. edulis* (Silva et al., 2019), vegetative maturation in *P. cincinnata* is also associated with pronounced changes in leaf morphology that occur gradually in successive phytomers and correlate negatively and positively with changes in miR156 and *SPL* transcript levels, respectively (Fig. S1). In *Arabidopsis*, *SPL*-dependent heteroblastic changes in leaf morphology are determined by altered rates of growth of different domains of the leaf and phase specific patterns on cell division and expansion. (Hu et al., 2023; Tang et al., 2023; Li et al., 2024). Whether similar mechanisms regulate progressively increased leaf lobing in *Passiflora* will be a key future area of research.

Constitutive expression of miR156 maintained *Passiflora* plants in the juvenile phase, supporting conservation of miR156 function across the angiosperms. However, a slight increase in leaf lobing was observed over time even in *PeOE* and *PcOE* plants (Fig. 2). This may reflect increased *SPL* transcription, as miR156 has been show to regulate *SPL* activity in a dose dependent manner (Fu et al., 2012; He et al., 2018). Conversely, it may reflect miR156-independent mechanisms of shoot maturation (He et al., 2018; Fouracre et al., 2021). Unlike the NT controls, neither *PeOE* nor *PcOE* plants flowered during the course of our experiments. This would suggest that *SPL* activity is critical for floral induction in *Passiflora,* at least in our growth conditions. Although miR156 does not play a major role in flowering time in *Arabidopsis* (Doody et al., 2022; Zhao et al., 2023; Poethig & Fouracre, 2024), *SPL* activity is an important determinant of flowering time in other species, including tobacco, alfalfa and switchgrass (Feng et al., 2016; Aung et al., 2015; Johnson et al., 2017).

### EFN formation is regulated in an age-dependent manner in *Passiflora*

Phase-specific traits, such as the presence of EFNs, have been previously identified during vegetative phase change in woody species (Leichty & Poethig, 2019; Machado et al., 2023). We showed here that the formation of EFNs is age-dependent in two *Passiflora* species and associated with heteroblastic changes to leaf morphology. During the juvenile vegetative phase, *P. edulis* produces simple leaves with petiolar EFNs, which are present consistently throughout vegetative development. During the transition from the juvenile to the adult vegetative phase, leaf morphology gradually changes and becomes tri-lobed (Silva et al., 2019), simultaneously forming nectaries on the inner side of the leaf margin furrow. EFNs are already present in the tri-lobed juvenile leaves of *P. cincinnata*; however, the increase in the number of leaf lobes leads to an increase in the abundance of nectaries at the margin, which was inhibited by high levels of miR156 (Figs. 3, 5). This observation indicates that *P. cincinnata* EFN formation/abundance in the leaf margin is at least partially dependent on the miR156/*SPL* module. Importantly, laminar EFN development and lobe formation increase concurrently, suggesting that these two aspects of leaf development are both regulated by an age-dependent pathway. Interestingly, a similar phenomenon is observed in leguminous *Vachellia* species, in which age-dependent production of EFNs evolved independently (Leicthy & Poethig, 2019). This would suggest that the miR156-*SPL* module has been coopted repeatedly during angiosperm evolution to regulate the timing of EFN production.

The size and abundance of laminar EFNs are greater in the adult phase, concomitant with an increase in leaf complexity and *SPL* transcript levels. In *Passiflora* miR156-OE lines, the size of the leaf lamina, as well as the size of petiolar and laminar EFNs, was reduced (Fig. 2), suggesting that the ontogeny of these structures was affected by low *SPL* expression and that changes in leaf development affect the growth of EFNs. Although the morphology of EFNs was affected in miR156-OE *Passiflora* plants, their tissue patterning was similar to NT plants, with demarcated nectary epidermis and nectariferous and subnectariferous parenchyma (Lemos et al., 2017; Moraes et al., 2022). However, cell number was strongly reduced mostly in the parenchymatic tissues (Fig.4m, n). These findings suggest that *SPL* genes regulate overall growth of EFNs, rather than specifying EFN cell identity. The effect of SPL proteins on rates of cell division is consistent with recent findings in *Arabidopsis* and tomato (Tang et al., 2023; Ferigolo et al., 2023; Li et al., 2024).

### miR156 regulates genes associated with *Passiflora* leaf patterning and EFN development

In addition to regulating leaf identity, the miR156/*SPL* module may also regulate EFN-associated genes. By evaluating the transcriptome data from *Passiflora* leaf primordia prior to EFN formation, we shed light on how miR156-targeted *SPLs* might regulate early EFN formation at the molecular level. Reduced *SPL* expression in miR156-OE leaf primordia led to the misexpression of genes associated with *Passiflora* leaf and EFN development, suggesting that these genes act together with the miR156-SPL module in a complex network (Nakayama et al., 2022). For instance, *AS1* transcript levels were slightly increased in miR156-OE plants (Fig. 5k; Tables S2, S3). Given that *AS1* is a member of the ARP family (*AS1*/ *RS2*/ *PHAN*) involved in adaxial identity and leaf complexity (Nakayama et al., 2022; Husbands et al., 2015), higher levels of *AS1* in miR156-OE may have affected the adaxial-abaxial patterning of leaf primordia, which is fundamental for proper EFN establishment.

Adaxial-abaxial polarity is also regulated by *YAB1* and *YAB5*. Importantly, *YAB5* is associated with the heteroblastic leaf development and floral nectary formation (Kram et al., 2009; Ostria-Gallardo et al., 2016; Chen et al., 2021). Thus, the presence of stunted EFNs in *Passiflora* miR156-OE leaves (Fig. 2) may be partially due to the low expression levels of *YAB1* and *YAB5* in the leaf primordia (Fig. 3; Table S2, S3). High levels of *LAX2-like* (Fig. 3; Table S2, S3), an auxin importer strongly associated with *Arabidopsis* leaf margin serration (Kasprzewska et al., 2015), might have led to impaired lobe formation in *Passiflora* miR156-OE leaf margins. Interestingly, tomato miR156-targeted *SPL15* negatively regulates auxin transport in the axillary buds partially via *LAX2* (Barrera-Rojas et al., 2023). Transcript levels of several *AGLs* were lower in miR156-OE (Fig. 3; Table S2, S3), suggesting that the activation of these genes may depend on *SPL* expression. The use of distinct *Arabidopsis* floral homeotic mutants demonstrated that proper floral nectary development requires *AG* and *AGL* genes (Slavković et al., 2021). It is possible, therefore, that a miR156-*SPL*-*AGL* genetic pathway is deployed to regulate both FN and EFN formation in *Passiflora* species.

### The changes in the morphology and sugar profile of miR156-OE EFNs affect Passiflora ecological relationships

EFNs promote mutualistic interactions with ants that act as defenders against herbivores in exchange for nectar, thereby establishing an indirect defense mechanism (Elias, 1983; Apple & Feener, 2001; Heil et al., 2015). Interestingly, EFNs increase in number in both *P. edulis* and *P. cincinnata* as the plant ages, suggesting that ant recruitment is prioritized during the adult phase. Although miR156-OE EFNs differentiate all the tissues necessary for nectar production and exudation, ant visitation was lower in miR156-OE lines of both *Passiflora* species. Changes in nectary size and abundance may partially explain the indifferent behavior of ants towards miR156-OE plants. Species with abundant, more prominent and accessible EFNs attract more ants than those with smaller nectaries, increasing defense performance (Apple & Feener, 2001; Alencar et al., 2023). Removing EFNs reduces ant patrolling, but mechanically damaging EFNs without affecting nectar supply has not been shown to limit ant visitation (Izarigue et al., 2013). Other variables, such as nectar quality and quantity, interfere with plant-animal mutualistic interactions (Heil et al., 2011; Alencar et al., 2023). The altered sugar profile in miR156-OE EFNs may have contributed to the observed reduction in ant patrolling. It is important to note that the lower sucrose content might be associated with lower expression of *SUC2-like* transporters in miR156-OE leaf primordia as, during the nectar secretion phase of wild type EFNs, genes encoding sucrose transporter are up-regulated (Chatt et al., 2021). Nectar with high concentrations of sugars seems to be more attractive to ants than those with low concentrations, resulting in increased visitation and persistence of the protective agent on the plant (Villamil et al., 2013; Alencar et al., 2023).

The nectar of EFNs in *Passiflora* species is chemically diverse and may contain alkaloids and flavonoids, and its consumption may depend on the metabolic capacity or preference of the consumer (Heil et al., 2011; Cardoso-Gustavson et al., 2013). Therefore, we cannot exclude the possibility of changes in the content of other chemical compounds present in the nectar of *Passiflora* miR156-OE EFNs may lead to a reduction in ant visitation. Because EFN visitors are defined as opportunistic species and do not always provide defensive assistance, it was not possible at this point to establish a specific mutualist relationship between the evaluated ants and *Passiflora* species. In addition, arthropod-plant interactions mediated by EFNs can be rapidly established between species that do not have a co-evolutionary relationship (Heil et al., 2015).

In summary, our observations indicate that the miR156-targeted *SPL* function may be fundamental for the establishment of EFNs in two *Passiflora* species with distinct leaf shapes. Further studies of molecular mechanisms that trigger extrafloral nectary establishment and patterning may provide valuable information for *Passiflora* yield improvement, as EFNs are key structures involved in strategies against herbivory in the genus *Passiflora*.

## Data availability

The RNA-Seq data underlying this article are available in Gene Expression Omnibus (GEO) of NCBI. All relevant data can be found within the article and its supplementary information, and they are openly available upon request.

## Supporting information

Supplementary Table S1. Oligonucleotide sequences used in this study.

Supplementary Table S2. Differentially Expressed Genes (DEGs) in leaf primordia of non-transgenic vs miR156-overexpressing Passiflora edulis.

Supplementary Table S3. Differentially Expressed Genes (DEGs) in leaf primordia of non-transgenic vs miR156-overexpressing Passiflora cincinnata.

Supplementary Table S4. Characteristics of SPL/SBPs genes in Passiflora species and the miR156 recognition region.

Supplementary Table S5. Putative miR156 targets in Passiflora edulis identified using the psRNATarget tool.

## Acknowledgements

We kindly thank Dr. Fabio G. Faleiro (EMBRAPA Cerrados, Brazil) and Viveiros Flora Brasil (Araguari, MG, Brazil) for donating the seeds. Dr. A. R. Leichty for kindly making available the miR156-overexpressing vector, and University of Pennsylvania for the infrastructure. We also thank the Universidade Federal de Viçosa (UFV) for providing all relevant infrastructure, Gilmar Valente from Núcleo de Microscopia e Microanálises (NMM) for SEM analyzes, and the Genética Molecular de Bactérias Laboratory (DMB – BIOAGRO) for RT-qPCR equipment.

## Author contributions

J.R.S., K.J.M.R., F.TS.N, and W.C.O. conceived and designed the study; W.C.O., J.R.S. and K.J.M.R performed the experiments; V.C.S. analyzed transcriptome data; L.L.L.D., L.A.S.S., E.C.S., C.S.S., P.A.S., provided technical assistance; Y.C.G.S identified the ant species; E.M.M., L.F.V. performed flow cytometry analysis; T.C.A.F. and E.R. analyzed the phylogeny data; L.S.F. and V.M.G. performed HPLC-based sugar analysis; F.G.P. performed the 5’RACE analysis. J.R.S., K.J.M.R., V.C.S., E.R., J.F, W.C.O. and F.T.S.N drafted and revised the manuscript.

## Supplementary Material

**Supplementary Figure 1.**
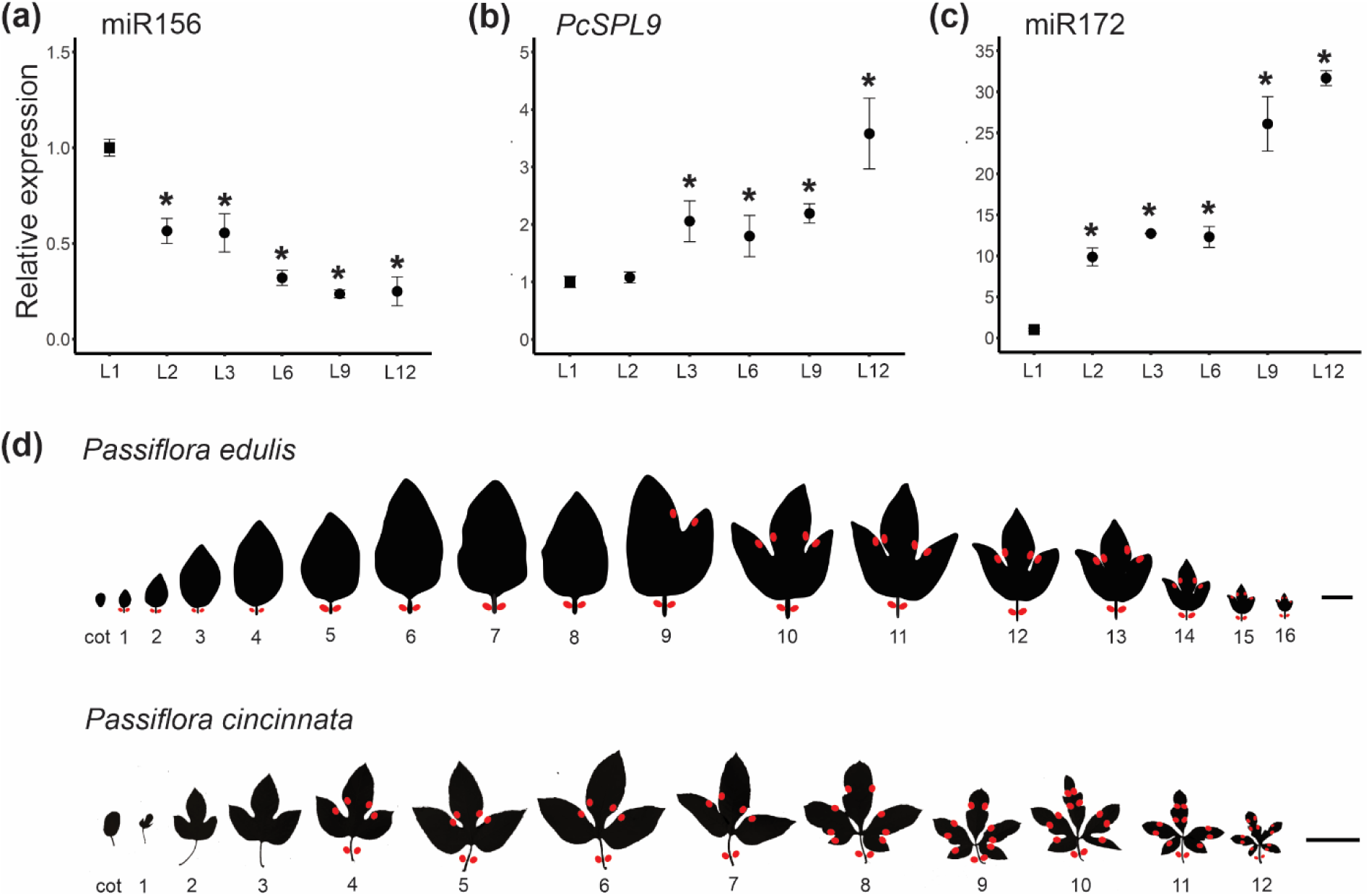
Expression patterns of microRNAs and *SPL9* during heteroblastic *Passiflora* leaf development. (a-c) Expression patterns of miR156 **(a)**, *PcSPL9* (b), and miR172 **(c)** in *Passiflora cincinnata* leaf primordia. Values are mean ± SE (n=3-4). \**P* ≤ 0.05 (Student’s *t*-test). **(d)** Overview of leaf heteroblastic development and location of extrafloral nectary (in red) in *Passiflora* species. Left to right, from young to adult leaves. Cot, cotyledons. Scale Bar: 2 cm; 10 cm.

**Supplementary Figure 2.**
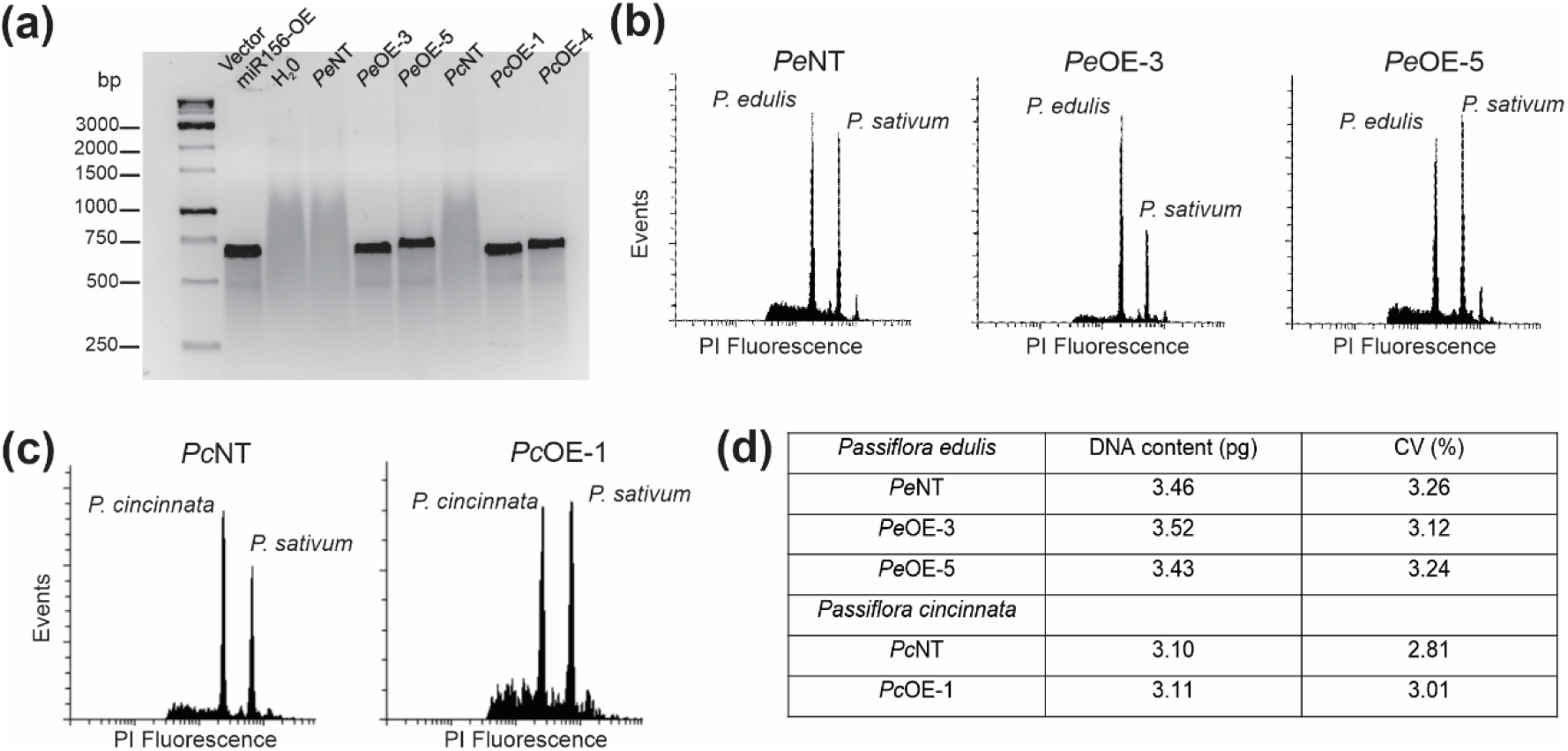
Transgenic molecular characterization and estimation of nuclear DNA content. **(a)** PCR amplification confirming the presence of the *hpt*II gene for hygromycin resistance in miR156-overexpressing plants (amplicon = 558 bp). **(b, c)** Flow cytometry analysis of non-transformed (*Pe*NT) and miR156-overexpressing events in *Passiflora edulis* (*Pe*OE-3 and *Pe*OE-5) **(b)**, and *Passiflora cincinnata* non-transformed (*Pc*NT) and miR156-overexpressing event (*Pc*OE-1) **(c)**. Each histogram first and second peaks correspond to the *Passiflora* and the *Pisum sativum* internal standard (2C = 9.09 pg of nuclear DNA). **(d)** Means of DNA content, determined by flow cytometry, in NT and miR156-overexpressing events. CV (%): Coefficient of variation.

**Supplementary Figure 3.**
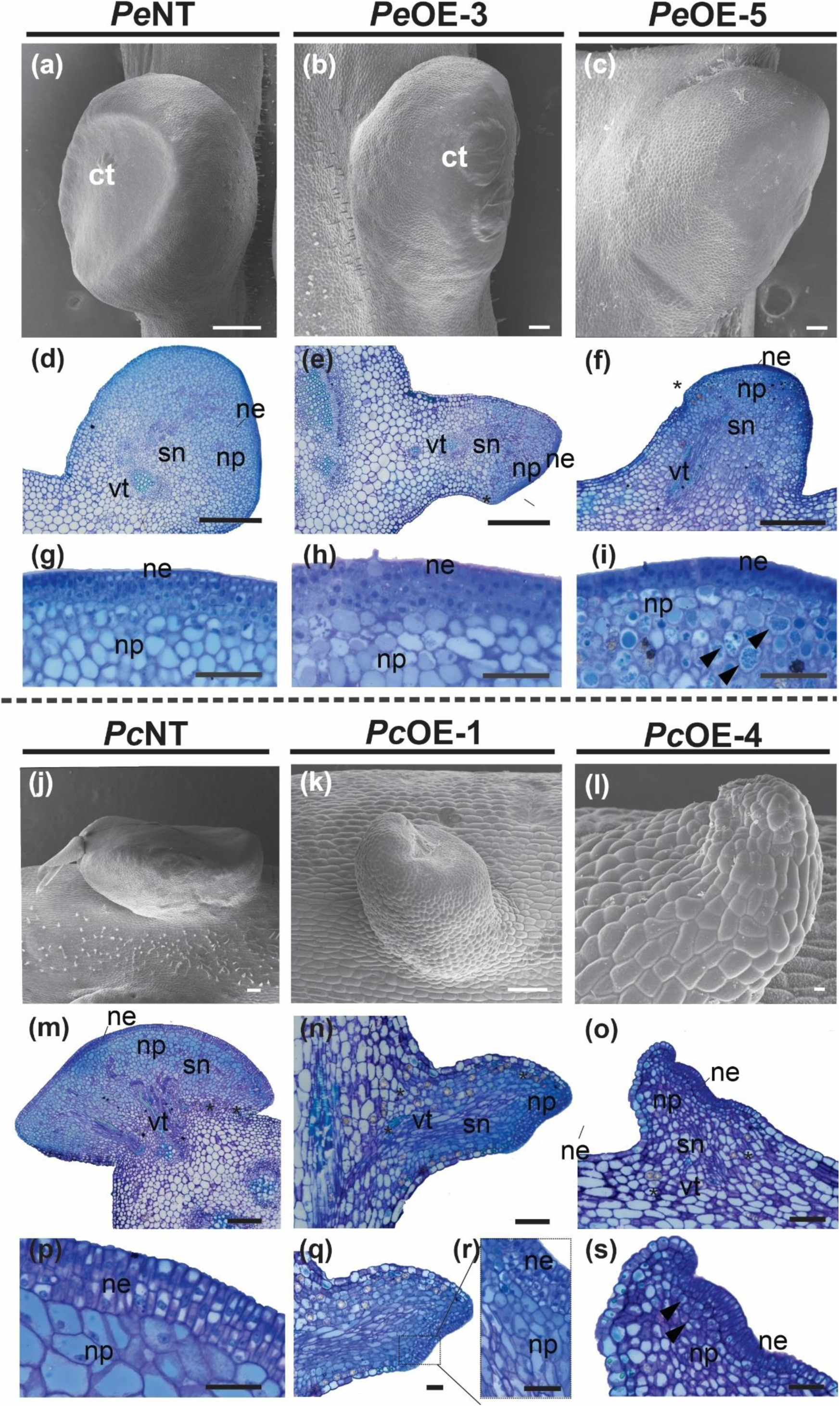
Morphology of *Passiflora* petiolar extrafloral nectaries (EFNs) is altered by the miR156 overexpression. Petiolar EFN Scanning electron microscopy (SEM) analysis in *Passiflora edulis* non-transgenic (*Pe*NT) **(a)**, *Pe*OE-3 **(b)** and *Pe*OE-5 **(c)** plants. Note the prominent cuticle with possible nectar content in the subcuticular region. Light microscopy *Passiflora edulis*: **(d, e)** Transverse sections of the leaf petiole showing petiolar EFN in *Pe*NT (**d**) and *Pe*OE-3 **(e). (f)** Longitudinal section in *Pe*OE-5 plants showing: Nectary epidermis (ne), nectariferous parenchyma (np); subnectariferous parenchyma (sn); leaf vascular tissue (vt) cells containing calcium oxalate crystals (*). (**g-i)** Detail of a petiolar EFN in *Pe*NT (**g**), *Pe*OE-3 **(h),** and *Pe*OE-5 **(i)**. **(j-l)** Petiolar EFN SEM in *P.cincinnata* in *Pc*NT(j), *Pc*OE-1 **(k),** and *Pc*OE-4 **(l)**. Note the ruptured cuticle in petiolar EFN *Pc*NT. **(m)** Transversal section of the petiole of leaf showing a petiolar EFN *Pc*NT. **(n,o)** Longitudinal sections of the *Pc*OE-1 **(n)** and *Pc*OE-4 **(o)** showing the three tissues of the nectary. **(p-r)** Detail of a petiolar EFN in *Pc*NT **(q)** and *Pc*OE-1 **(r)**, highlighting multiseriated epidermis. **(s)** Detail of a petiolar EFN in *Pc*OE-4, Arrowheads showing dense cellular content. Scale bar: a = 200 μm, b-c = 10 μm, d-f = 150 μm, g-i =50 μm, j-k = 100 μm, l = 10 μm, m-o= 200 μm, p= 50 μm, q-s=100 μm.

**Supplementary Figure 4.**
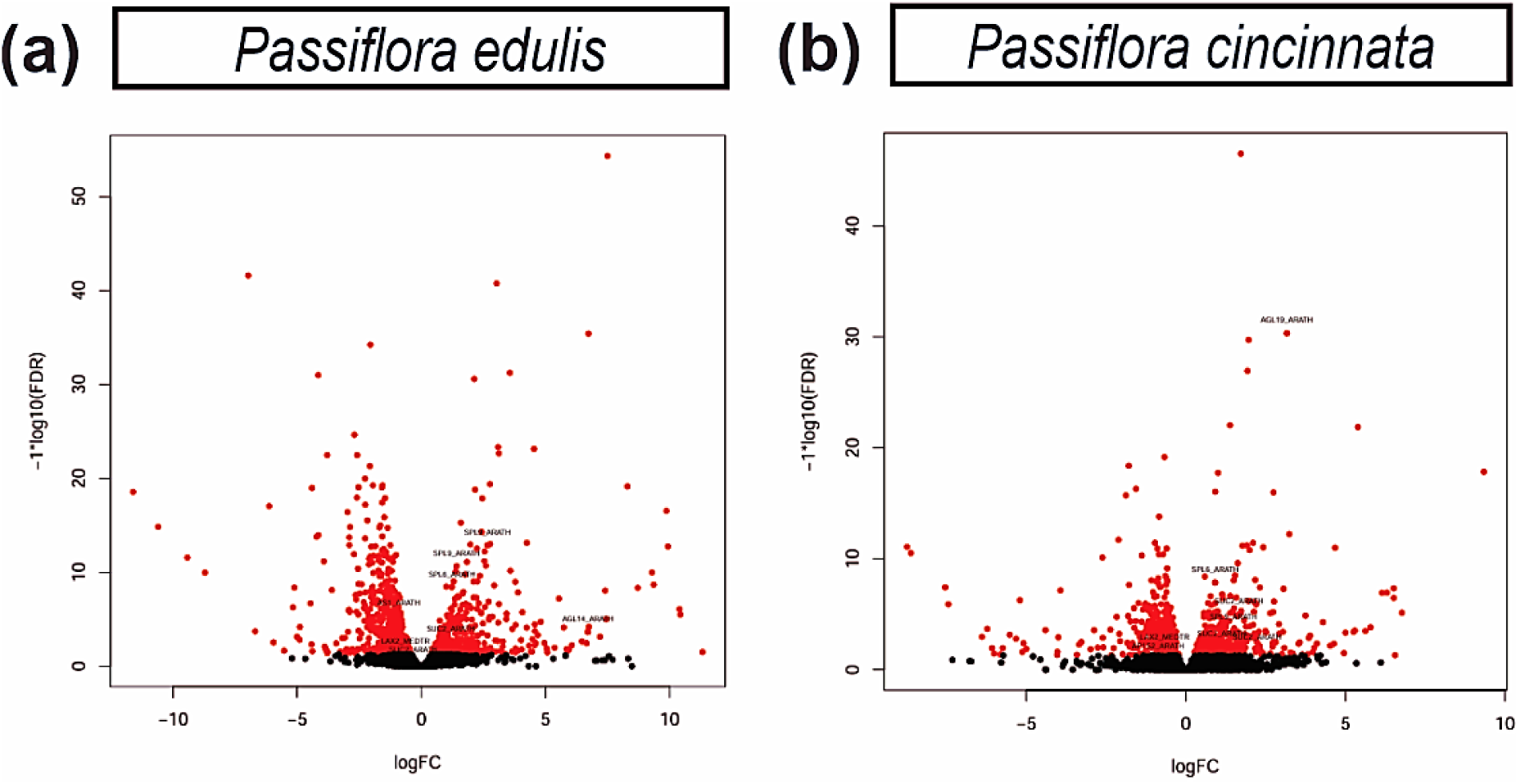
Distribution of differentially expressed genes (DEGs) from the RNA-seq data. (a, b) Volcano plot of DEGs representing up-(LogFC >0) and down-regulated (LogFC<0) genes in red color from non-transgenic vs miR156-overexpressing *Passiflora edulis* **(a)** and *P. cincinnata* (**b**) leaf primordia.

**Supplementary Figure 5.**
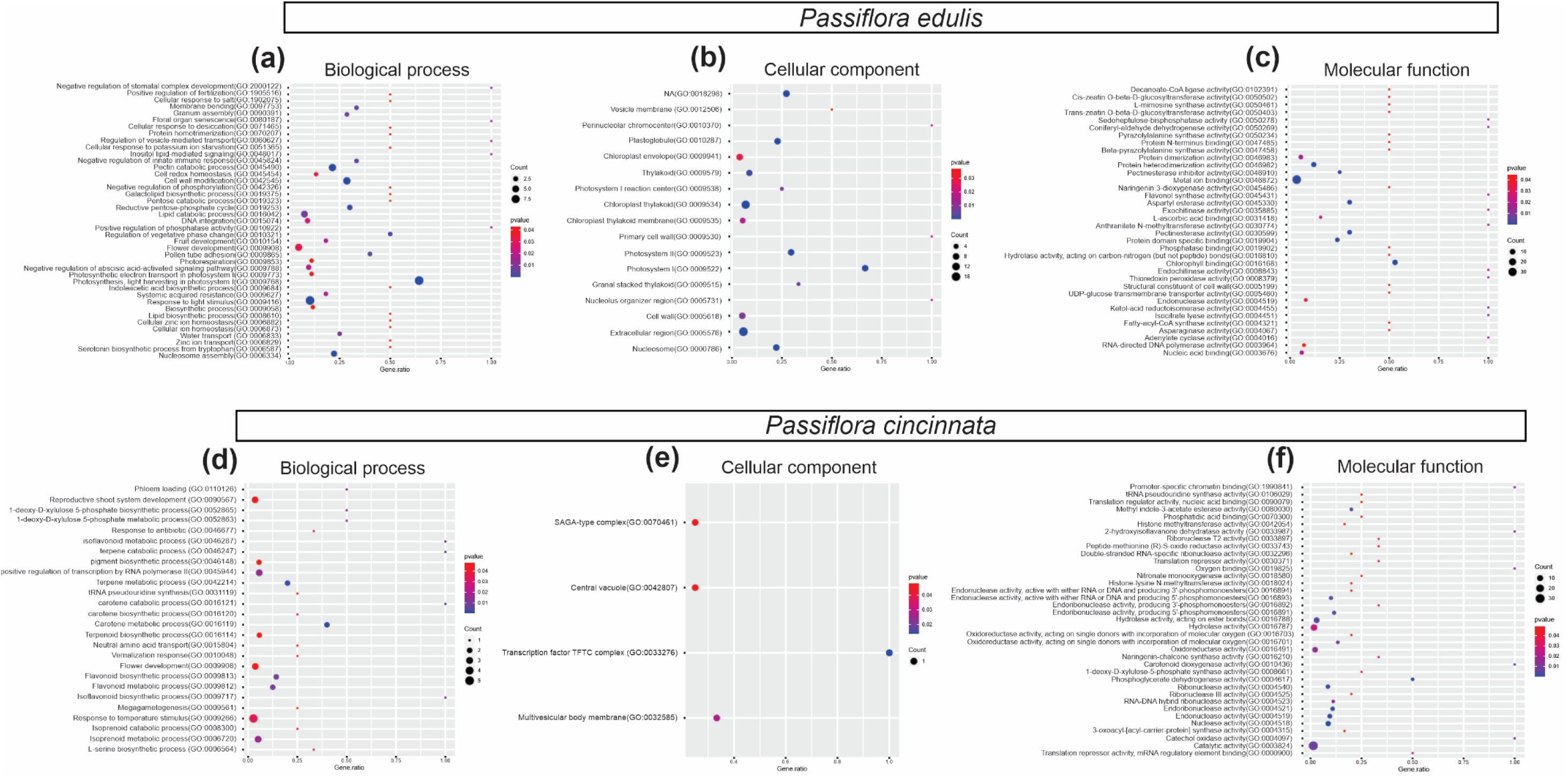
Gene Ontology enrichment of DEGs in miR156-overexpressing versus non-transgenic plants. **(a)** Biological process **(b)** cellular component and **(c)** Molecular function in *Passiflora edulis*. **(d)** Biological process **(e)** cellular component and **(f)** Molecular function in *Passiflora cincinnata*. The dot size represents the number of DEGs associated with the process (count) and the dot color represents the significance of the enrichment (p-value ≤ 0.05).

**Supplementary Figure 6.**
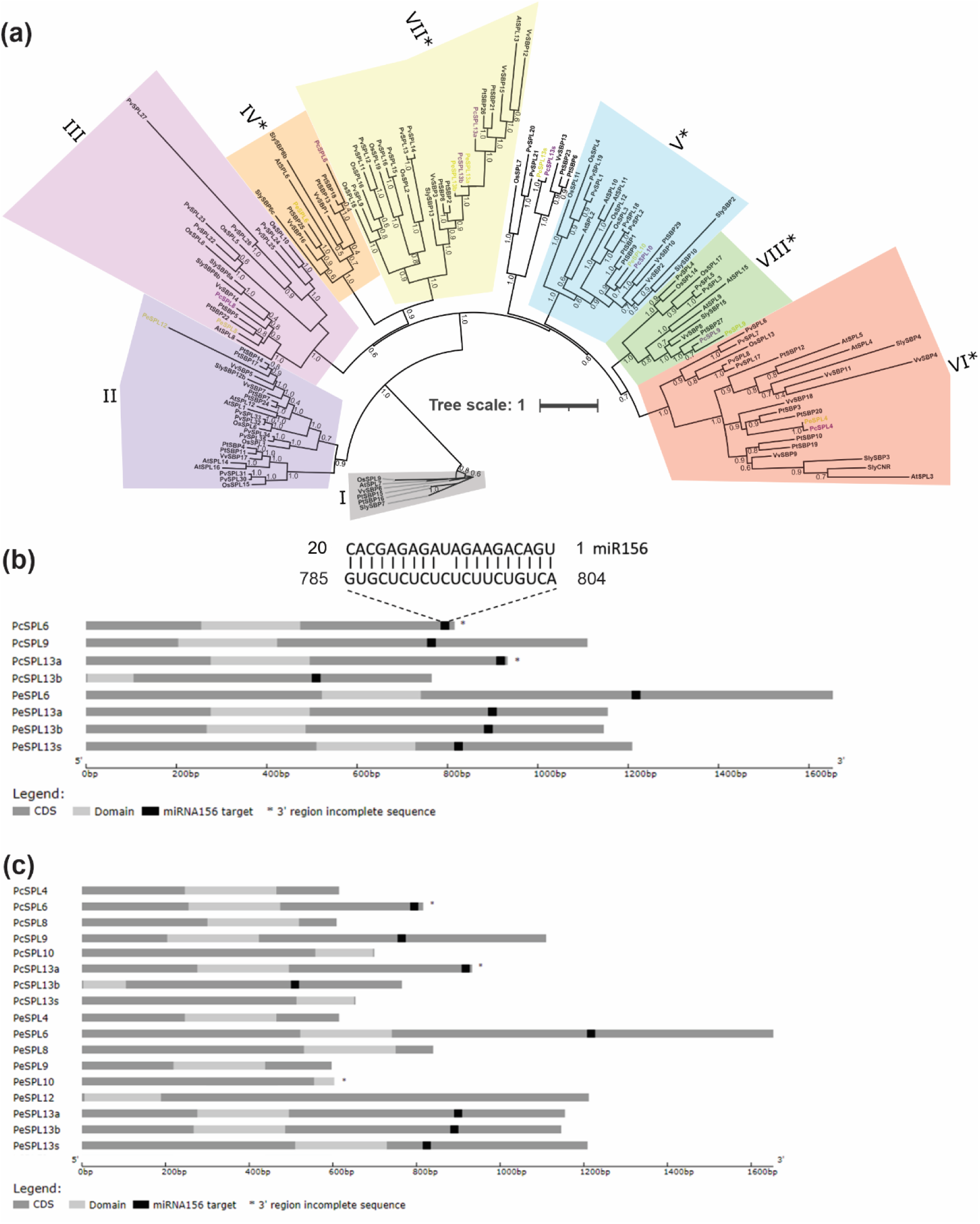
Maximum-likelihood phylogeny of the miR156- targeted SPL/SBP proteins of *Passiflora* species. **(a)** Unrooted phylogenetic tree based on an alignment of all SPL/SBP sequence encoded protein obtained for *P. cincinnata* (*PcSPL*), *P. edulis* (*PeSPL*), tomato (*SlySBP*), Arabidopsis (*AtSPL*), poplar (*PtSBP*), grapes (*VvSBP*), rice (*OsSPL*), and swithgrass (*PvSPL*), using MUSCLE tool and Interactive Tree of Life (iTOL) resource to annotate. Original code gene for each species is available (Table S4). Clade colors match Salinas et al. (2012) where applicable. Clade with asterisks is regulated by miR156 or miR157, as stated by Preston and Hileman (2013). The code names of *P. edulis* and *P. cincinnata SPLs* were marked in yellow and purple, respectively, for highlight; those with asterisks contain predictable target region for miR156 in their sequence (Table S5). Scale bars indicate the number of substitutions per site. **(b)** Heavy grey lines represent ORFs. The light grey box represents SBP domain. miRNA complementary sites (black box) with the nucleotide positions of *Passiflora SPLs* sequences are indicated. Asterisk means incomplete 3’region from a detected transcript. **(c)** *Passiflora SPLs* non-targeted and targeted by miR156. Heavy grey lines represent ORFs. The light grey box represents SBP domain. miRNA complementary sites (black box) with the nucleotide positions of *Passiflora SPLs* sequences are indicated. Asterisks mean incomplete 3’ sequences.

**Supplementary Figure 7.**
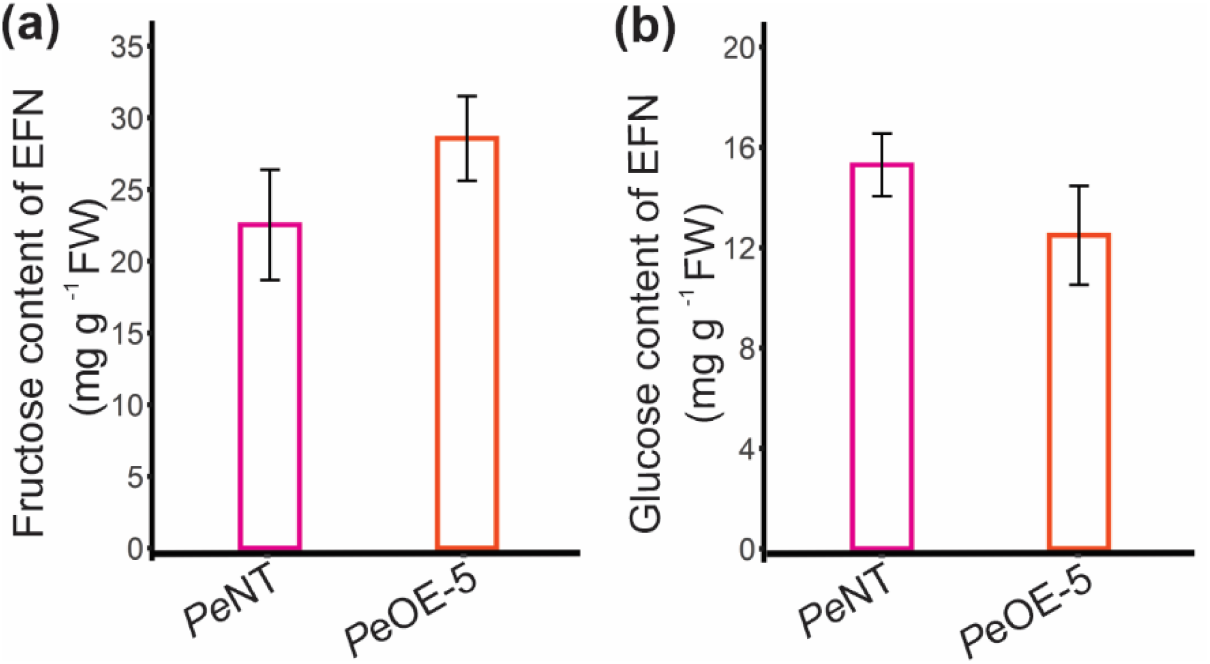
Sugar profiles in miR156-overexpressing *Passiflora edulis* extrafloral nectaries (EFNs). **(a)** Fructose and **(b)** glucose content of the laminar and petiolar extrafloral nectary pool of *Passiflora edulis* non-transgenic (*Pe*NT) and miR156 overexpressor (*Pc*OE-5) (n= 3).

**Supplementary Table S1.** Oligonucleotide sequences used in this study.

**Supplementary Table S2.** Differentially Expressed Genes (DEGs) in leaf primordia of non- transgenic vs miR156-overexpressing *Passiflora edulis*.

**Supplementary Table S3.** Differentially Expressed Genes (DEGs) in leaf primordia of non- transgenic vs miR156-overexpressing *Passiflora cincinnata*.

**Supplementary Table S4.** Characteristics of *SPL/SBPs* genes in *Passiflora* species and the miR156 recognition region.

**Supplementary Table S5.** Putative miR156 targets in *Passiflora edulis* identified using the psRNATarget tool.

